# ISG15-modification of the Arp2/3 complex restricts pathogen spread

**DOI:** 10.1101/2022.12.27.522022

**Authors:** Yifeng Zhang, Brittany Ripley, Wei Ouyang, Miranda Sturtz, Ellen Upton, Emma Luhmann, Madeleine Vessely, Rocio Coloma, Nathan Schwery, Scott M. Anthony, Adam Goeken, Thomas O. Moninger, John T. Harty, Aloysius Klingelhutz, Emma Lundberg, David K. Meyerholz, Balaji Manicassamy, Christopher Stipp, Susana Guerra, Lilliana Radoshevich

**Affiliations:** Department of Microbiology and Immunology, University of Iowa Carver College of Medicine, Iowa City, IA, 52242, USA; Science for Life Laboratory, School of Engineering Sciences in Chemistry, Biotechnology and Health, KTH – Royal Institute of Technology, Stockholm, Sweden; Department of Bioengineering, Stanford University, Stanford, CA, 94305, USA; Department of Preventive Medicine, Public Health and Microbiology, Universidad Autónoma, E-28029 Madrid, Spain; Department of Pathology, University of Iowa Carver College of Medicine, Iowa City, IA, 52242, USA; Central Microscopy Research Facility, University of Iowa Carver College of Medicine, Iowa City, IA, 52242, USA; Chan Zuckerberg Biohub, San Francisco, San Francisco, CA, 94158, USA; Department of Biology, University of Iowa, Iowa City, IA, 52242, USA

**Keywords:** ISG15, ubiquitin-like proteins, Arp2/3, host-pathogen interactions, cell motility, cell-to-cell spread, Interferon, *Listeria monocytogenes*, Vaccinia virus

## Abstract

The ubiquitin-like protein, ISG15, can act as a cytokine or can covalently modify host and pathogen-derived proteins. The consequences of ISG15 modification on substrate fate remain unknown. Here we reveal that ISGylation of the Arp2/3 complex slows actin filament formation and stabilizes Arp2/3 dependent structures including cortical actin and lamella. When properly controlled, this serves as an antibacterial and antiviral host defense strategy to directly restrict actin-mediated pathogen spread. However, *Listeria monocytogenes* takes advantage in models of dysregulated ISGylation, leading to increased mortality due to augmented spread. The underlying molecular mechanism responsible for the ISG15-dependent impact on actin-based motility is due to failed bacterial separation after division. This promotes spread by enabling the formation of multi-headed bacterial “bazookas” with stabilized comet tails that can disseminate deeper into tissues. A bacterial mutant that cannot recruit Arp2/3 or a non-ISGylatable mutant of Arp3 is sufficient to rescue slowed comet tail speed and restrict spread. Importantly, ISG15-deficient neonatal mice have aberrant epidermal epithelia characterized by keratinocytes with diffuse cortical actin, which could underlie observed defects in wound healing in human patients who lack ISG15. Ultimately, our discovery links host innate immune responses to cytoskeletal dynamics with therapeutic implications for viral infection and metastasis.

## Main text

The Interferon stimulated gene 15 (ISG15) is an innate immune effector that exhibits broad and potent antimicrobial activity^1^. ISG15 can be secreted and act as a cytokine^2^, can non-covalently contribute to antiviral responses^3^, and can covalently modify host and microbial proteins through a ubiquitin-like enzymatic cascade^4–7^. This process, called ISGylation, is reversible and deconjugation is mediated by an ISG15-specific isopeptidase (USP18)^8^. ISG15 is mutated in a cohort of patients, who are susceptible to Mycobacterial diseases^2^, have Aicardi-Goutières-like Interferonopathies^9^, and non-resolving skin lesions^10,11^. These patients brought to light the role of free ISG15 as a cytokine^12^ and in intracellular JAK/STAT signaling^9^, however the consequences of covalent ISG15 modification remain poorly understood despite perturbation of ISGylation in important human pathologies such as cancer and neurodegenerative diseases.

Type I Interferon is almost universally antiviral^13^, however paradoxically, some bacterial pathogens directly^14^ or indirectly^15–17^ elicit ISG expression to promote replication, spread, and to reduce T-cell mediated immunity^18^. In particular, the Gram-positive food borne pathogen *Listeria monocytogenes* (*Lm*)^19^ requires Type I IFN signaling for infection^20–22^. ISG15 acts as an antibacterial effector in a conjugation-dependent manner if properly regulated^23,24^, however if its isopeptidase, USP18, is replaced with a catalytically inactive mutant^25^, *Lm* can take advantage exhibiting increased spread in the liver^26^. Since ISG15 covalently modifies target proteins and the consequences of conjugation on protein fate remains elusive, we explored the mechanism of enhanced *Lm* spread under conditions of enhanced ISGylation.

Here we uncover a novel role for ISG15 modification of the ARP2/3 complex in restriction of microbial spread. ISG15 covalently modifies ARP3 following infection, which alters *Lm* and Vaccinia virus (VacV) actin-comet tail morphology. Unlike Vaccinia virus, *Lm* takes advantage by dividing without disassociating from the stabilized comet tail ultimately resulting in enhanced spread and increased mortality following *Lm* infection. Furthermore, ISG15-stabilized ARP2/3 occurs in cortical actin and slows cell motility, increases resistance to trypsin, and alters epithelial barrier integrity in neonatal mice, a mechanism which could underlie the phenotype in human patients with non-resolving skin lesions.

### ISG15 affects bacterial cell-to-cell spread

To dissect the functional consequences of ISGylation in *Lm* infection, we infected *usp18*^+/+^ (WT) and *usp18^C61A/C61A^* (CA) mutant mice. The CA mutant of USP18 is catalytically inactive, thus infection of these mice results in a state of enhanced ISGylation. In contrast to VacV infection in which CA is protective^25^, CA mice all died between Day 4 and Day 7 following *Lm* infection (Fig.1a); whereas all WT mice survived despite negligible differences in weight loss (Extended Data Fig.1a). In order to assess bacterial spread in vivo we performed intravital microscopy at three days post infection (Fig.1b, Extended Data Fig. 1b-c) and observed that bacteria in WT animals were primarily contained within phagocytic cells (F4/80+) whereas in CA mice bacteria also localized to hepatocytes (F4/80-) (Fig.1c). These findings corroborate our data from histological sections of infected liver^26^ (Extended Data Fig.1b-c) and suggest that enhanced ISGylation promotes bacterial cell-to-cell spread. Using a bacterial plaque assay, we subsequently infected confluent monolayers of WT, CA, or *isg15^−/−^* (KO) mouse embryonic fibroblasts (MEFs) to determine whether there is a difference in bacterial cell-to-cell spread in the absence of immune cells. Indeed, at three days post infection *Lm* formed larger plaques in both CA and KO cells than in WT cells, which recapitulates the increased bacterial focal area we observe in vivo (Fig.1d-e). Therefore, while initially protective, unchecked ISGylation ultimately favors bacterial spread both in vitro and in vivo.

**Figure 1:**
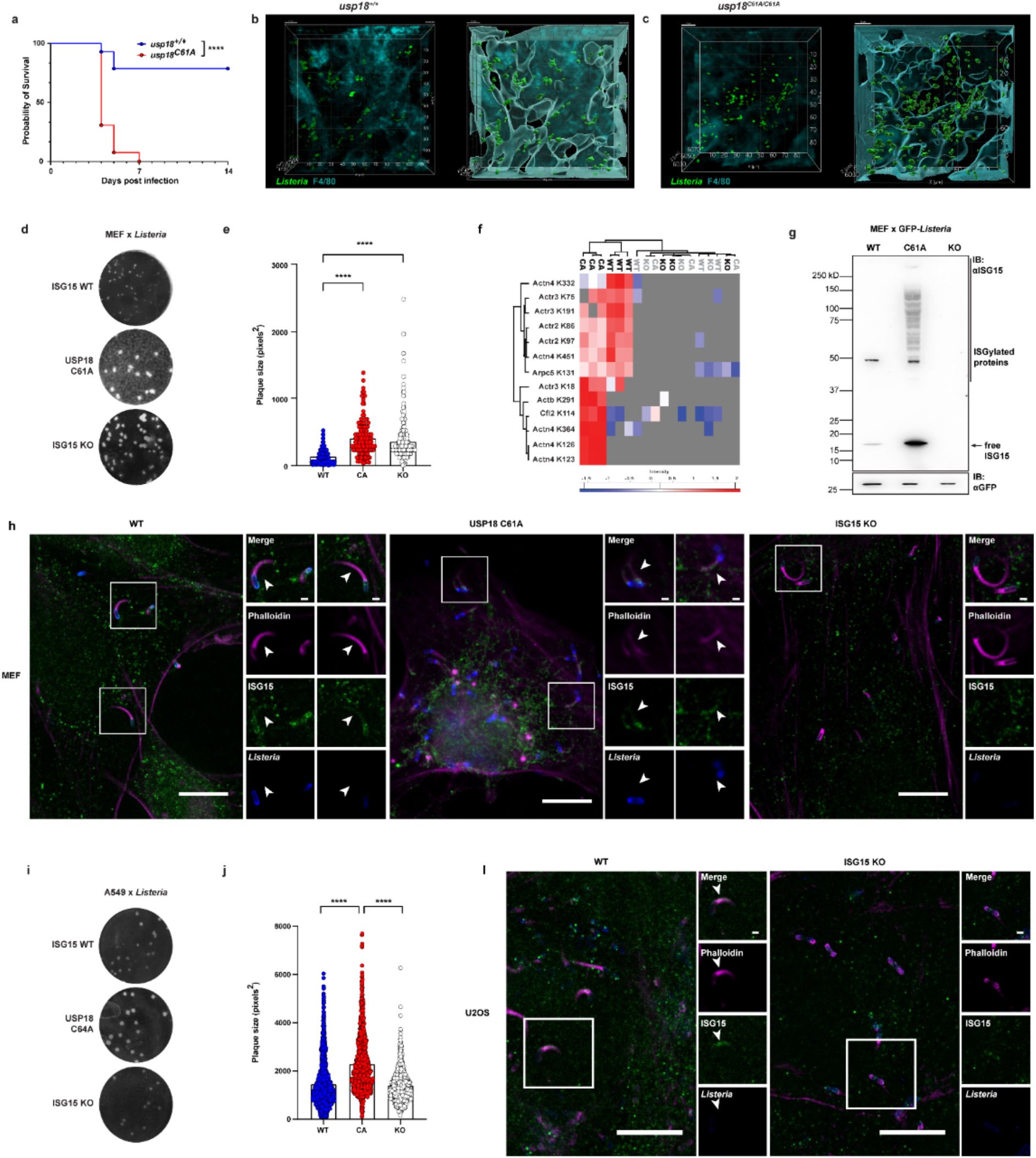
ISG15 modifies actin-comet tail proteins and affects *L. monocytogenes* cell-to-cell spread. **a** Survival data from C57BL6/J mice (WT and *usp18^C61A/C61A^*) infected intravenously with 1 × 10^4^ Colony Forming Units (CFUs) of *Listeria monocytogenes* (*Lm*) strain 10403s; results are from two individual repeats, wild-type n = 14, USP18^C61A/C61A^ n = 13, significance determined using Log-rank test. **b-c** Two-photon microscopy from livers of live WT (**b**) and *usp18^C61A/C61A^* (**c**) mice infected with 5 × 10^5^ CFUs of *Lm* GFP-EGD (in green) and F40/80 positive cells (in teal) 3 days post infection (DPI). Microscope image (left) and 3D rendering (right) Scale bar, 10 μm. **d** Representative image of plaque formation in Mouse Embryonic Fibroblasts (MEFs) infected with *Lm* EGD using a multiplicity of infection (MOI) of 0.05 in WT, USP18^C61A/C61A^ and *isg15^−/−^* cells. **e** Measurement of plaque size (in square pixels) from three independent experiments (WT n=202, CA n=128, KO n=422, statistical analysis was conducted using Kruskal-Wallis test with Dunn’s postdoc). **f** Heatmap showing significantly enriched lysine sites identified on actin-comet tail proteins following *Lm* infection. **g** SDS-PAGE analysis of GFP-EGD *Lm* purified from WT, USP18^C61A/C61A^, and *isg15^−/−^* MEFs (MOI 10) at 24 HPI. **h** Representative images of infected MEFs (*Lm* EGD MOI of 10) for 6h. anti-ISG15 depicted in green, actin (phalloidin) in magenta, and bacteria (anti-ActA) in blue; scale bars in image and insets are 10 μm and 1 μm, respectively. White arrows point to colocalization. **i** Representative image of plaque formation of *Lm* 10403s infection (MOI 0.001) in WT, USP18^C64A/C64A^ and *isg15^−/−^* human lung epithelial cells (A549) 3 DPI. **j** Measurement of plaque size (in square pixels); data from three independent experiments (WT n=202, CA n=128, KO n=422), analyzed with Kruskal-Wallis test followed by Dunn’s post hoc test. **i**, Representative image of wildtype and *isg15^−/−^* human osteosarcoma cells (U2OS) infected with *Lm* EGD for 18 h; anti-ISG15 depicted in green, actin (Phalloidin) in magenta, and bacteria (a-ActA) in blue. Scale bars in the picture and insets are 10 μm and 1 μm, respectively. White arrows point to colocalization. In **a**, **e**, and **j** asterisks indicate p values with ****p < 0.0001. Raw data are available in the source data file.

Since increased plaque size indicates an effect of ISG15 on cell-to-cell spread, we hypothesized that modulators of actin dynamics could be modified by ISG15. We cross referenced a proteome of in vitro reconstituted *Listeria* actin comet tails^27^ to the endogenous ISGylome^26^ which we previously mapped in the liver of *Lm* infected WT and CA mice. We were able to identify thirteen ISG15 sites on six actin binding proteins, including three members of the Arp2/3 protein complex (Arp2, Arp3, and ArpC5), an essential actin nucleator, an actin filament severer (cofilin-2), an actin bundler (alpha-actinin 4), and monomeric actin itself (β-actin). While actin^28^ and alpha-actinin^23^ had been previously identified as targets of ISG15, this is the first report, to our knowledge, that the Arp2/3 complex and cofilin are ISGylated. Notably, these specific lysine sites only occur following *Lm* infection and were not detected to be modified by ubiquitin in the KO mice (Fig.1f). Furthermore, specific sites on Arp3, cofilin, alpha-actinin, and actin are solely modified in conditions of enhanced ISGylation. To validate our proteomics data and assess whether ISG15 modifies comet tails, we isolated bacteria from infected MEFs and visualized comet tail proteins using SDS-PAGE. Both free ISG15 and ISGylated proteins associate with *Lm* comet tails in WT and CA cells (Fig.1g). As predicted from our proteomics analysis, bacteria-associated proteins showed increased ISGylation in CA MEFs. Since our biochemical isolation could not differentiate between bacterial surface-associated or actin comet tail-localized ISG15 and ISGylation, we also imaged *Lm* in the cytosol of MEFs six hours post infection using immunofluorescence microscopy. In so doing, we could visualize ISG15 localized to *Lm* actin comet tails in WT and CA but absent in KO cells (Fig.1h). Taken together, this suggests that ISGylation occurs on the actin comet tail and is enhanced in isopeptidase-dead cells as predicted by our proteomic analysis.

Unlike ubiquitin which is highly conserved from yeast to man, human and mouse ISG15 only share sixty-three percent sequence identity. As a result, not all functions of ISG15 are conserved. We therefore interrogated whether enhanced ISGylation also promotes bacterial spread in human cells that either lack ISG15 or express an isopeptidase-dead CA mutant of USP18(C64A); the catalytic cysteine in human USP18 is cysteine sixty-four (Extended Data Fig.1d-e). *Lm* formed larger plaques in human epithelial cells with enhanced ISGylation compared to WT A549s and ISG15-deficient cells (Fig.1i-j, Extended Data Fig.1f-g). Interestingly, we no longer observed an increase in spread in human ISG15-deficient cells using a wildtype *Lm* strain 10430s, suggesting that enhanced spread in ISG15-deficient cells could depend on virulence factor expression levels which are higher in the EGD strain^29^ (Fig.1i-j). Using immunofluorescence microscopy, endogenous ISG15 localizes to *Listeria* actin comet tails in WT U2OS cells but not in KO cells as expected (Fig.1l, Extended Data Fig.1d). Taken together, these results establish that ISG15 modifies proteins in actin-comet tails both in mouse and human cells following *Lm* infection.

### ISGylation alters Arp3 localization on the bacterial comet tail

In order to determine the consequences of ISGylation on cell-to-cell spread, we assessed whether ISG15 modification affects the localization of target actin-modulators in *Lm* comet tails. In conditions of enhanced ISGylation, Arp2 and Arp3 aberrantly localize to the bacterial surface. Instead of decorating the actin comet tail as is the case for WT and KO cells, Arp3 also coats the bacterial surface (Fig.2a-c, Extended Data Fig.2a). When quantified, there were significantly (p=0.0096, p=0.0087) more bacteria in CA cells with Arp3 colocalization of the bacterial surface than in either WT or KO cells (Fig.2d). Interestingly, ISG15 modification did not appear to affect the surface or comet-tail localization of ArpC5, cofilin, or alpha-actinin (Extended Data Fig.2b-c, 2f). In order to better resolve the morphology of the actin comet tail we made use of stimulated emission depletion (STED) super-resolution microscopy. Ultra-structurally, at six hours post-infection actin, like Arp3, more frequently colocalized on the sides of bacteria in CA MEFs, though we also observe surface colocalization in some comets from WT MEFs (Fig.2e). Cells with enhanced ISGylation also harbor shorter tails than in WT or ISG15-deficient cells. In order to quantify these data in an unbiased manner we worked to establish a machine-learning model to identify actin comet tails to measure their morphological properties. In CA MEFs, *Lm* forms significantly shorter (p<0.0001) and thicker (p=0.0002) tails (Fig.2f-g), with stronger local fluorescence intensity (Fig.2h), corroborating that enhanced ISGylation shortens actin filament structure. We also employed STED super-resolution microscopy at twelve hours post infection, which revealed extremely striking actin structures in the CA mutant (Fig.1i). Actin colocalized with the surface of aberrantly elongated bacteria. Connected daughter bacteria with two stable tail structures occur in CA (Fig.1i, panel 1 and 3) but were not observed in WT or KO. Thus, enhanced ISGylation of the bacterial comet tail results in shorter actin structures which colocalize with the bacterial surface.

**Figure 2:**
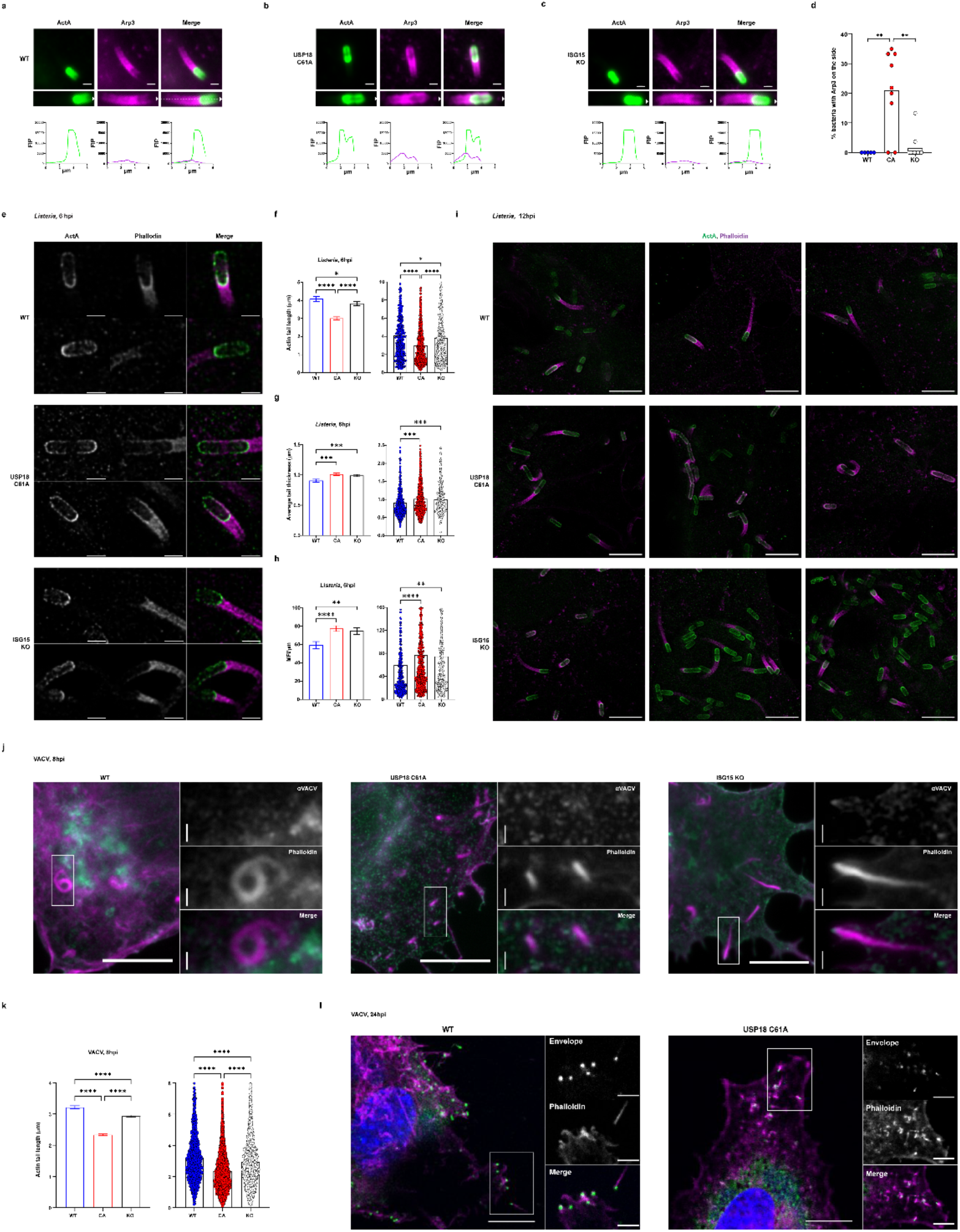
ISGylation affects Arp2/3 localization on the bacterial surface and length and intensity of comet tails in *Lm* and Vaccinia infection. **a-c** Arp3 actin comet tail in *Lm* EGD infected MEFs at 6 h post infection (MOI 10). Bacteria are shown in green, and Arp3 in magenta. Fluorescent intensity profile (FIP) was taken of the dotted line along the midline of the cell in the inset image WT (**a**), USP18^C61A/C61A^ (**b**), and *isg15^−/−^* (**c**), scale bar, 1 μm. **d** Percentage of bacteria with Arp3 surface localization at 6 HPI, representative data (WT n=5, CA n=9, KO n=10) analyzed with Kruskal-Wallis test followed by Dunn’s post hoc test. **e** Stimulated Emission Depletion (STED) images of bacteria in MEFs at 6 h post infection, bacteria are shown in green (a-ActA), and actin (Phalloidin) in magenta. scale bar, 1 μm. **f** Data from machine learning analysis of average length of actin comet tails formed by bacteria at 6 h post infection in wildtype, USP18^C61A/C61A^, or *isg15^−/−^* MEFs analyzed by Kruskal-Wallis test with Dunn’s post hoc. **g** Average thickness of actin comet tails formed by bacteria at 6 h post infection in wildtype, USP18^C61A/C61A^, or *isg15^−/−^* MEFs, compared by by Kruskal-Wallis test with Dunn’s post hoc. **h** Average signal intensity of actin structures along bacterial actin tails in wildtype, USP18^C61A/C61A^, or *isg15^−/−^* MEFs analyzed by Kruskal-Wallis test with Dunn’s post hoc. In **f-g** bar graphs represent the mean ± SEM and results analyzed from three independent experiments WT n=439, CA n=661, KO n=680. **i** Representative images of *Lm* actin comet tail formed at 24 HPI using STED imaging, bacteria in green (a-ActA), and actin (Phalloidin) in magenta; scale bar, 5 μm. **j** Wildtype, USP18^C61A/C61A^, and *isg15^−/−^* MEFs infected with Western Reserve strain of Vaccinia virus (VACV) for 8 h. Actin is shown in magenta, and VACV in green; scale bar,10 μm, insets, 1 μm. **k** Average length of VACV actin tail in MEFs; data represent the mean ± SEM, WT n=1183, CA n=3286, KO n=1752, significance determined by Kruskal-Wallis test with Dunn’s post hoc. **l** Representative confocal images of VACV actin comet tails at 12 HPI in WT and USP18^C61A/C61A^ using anti-anti-VACV 14K antibody (green) and phalloidin for actin (magenta); scale bar, 10 μm, insets, 1 μm. In **d**, **f**, **g**, **h** and **k** asterisks indicate p values with *p < 0.05, **p < 0.01, ***p < 0.001, and ****p < 0.0001. Raw data are available in the source data file.

Since *Lm* is one of several pathogens that co-opts the actin cytoskeleton to enable spread^30^, we next determined whether ISGylation of the Arp2/3 complex could be broadly antimicrobial by shortening and stabilizing viral actin tails. VacV can also form actin comet tails^31^ within the cell which emanate from cell-associated enveloped virions (CEV)^32^. Here we assessed actin tail length at eight and twenty-four hours post infection using the Western Reserve strain of Vaccinia virus. Indeed, as was the case for *Lm* we found that VacV formed significantly (p<0.0001) shorter actin comet tails in CA cells at 8 hours post infection (Fig.2k). At this time in the KO cells the tails were slightly longer than in CA cells, though still significantly (p < 0.0001) shorter than WT actin tails. Interestingly, at 24 hours post infection the phenotype is more pronounced with extremely short actin tails in CA. Since CEV spread on the surface of cells, these shortened tails could cause a block or delay in VacV dissemination (Fig.2l). CA cells and mice are highly resistant to VacV^25^. We posit that ISGylation of this complex restricts comet length, which leads to protection against VacV since Arp2/3 mediated actin tails are critical for effective viral spread. More importantly, our data indicate that ISGylation of mediators of actin-based motility acts a critical host defense strategy for both bacterial and viral pathogens, when properly regulated.

### ISG15 slows nucleation of actin comet tails by the Arp2/3 complex

The Arp2/3 complex turns over as the actin filament forms at the bacterial pole creating the force necessary to push bacteria forward^33,34^. Thus, we hypothesized that ISGylation of the Arp2/3 complex could affect bacterial speed since Arp3 aberrantly colocalized with the bacterial surface. When we compared bacterial motility across all three genotypes at four hours post infection, in CA cells *Lm* moves significantly more slowly than in WT or KO cells (p=0.0027, p=0.0002). On average *Lm* moves through WT MEFs at 0.04 μm/second (2.4 μm/minute), whereas in CA cells the average speed slowed to 0.017 μm/second (1.02 μm/minute), while the KO cells housed the fastest moving bacteria at 0.068 μm/second (4.08 μm/minute) (Fig.3a-b). Interestingly, in cells with enhanced ISGylation we observed instances of bacteria either being unable to detach from a thick actin cocoon-like structure (SI movie 2-3) or having an initially motile bacterium where the tail stops treadmilling and covers the surface of *Lm* similar to the colocalization we observed in fixed microscopy (SI movie 2, Fig.3a). On the other hand, bacteria moved visibly faster in KO cells though sometimes on a circuitous path (Fig.3a-b). We quantified straightness of the track and in the ISG15-deficient cells the bacteria were much more likely to turn relative to the path in cells with enhanced ISGylation (Extended Data Fig.2d-e). In KO MEFs actin comet tails were longer and persist resulting in residual actin tracks which trace where the bacteria have been within the cell. Our machine-learning analysis could not identify these long tails by fixed microscopy (Fig.2h-j), likely because breaks in the tail due to changed depth in the z plane or collision with the edge of the cell would result in measuring the length of only part of the tail. Taken together, enhanced ISGylation slows the speed of actin filament assembly whereas the lack of ISG15 increased assembly speed.

**Figure 3:**
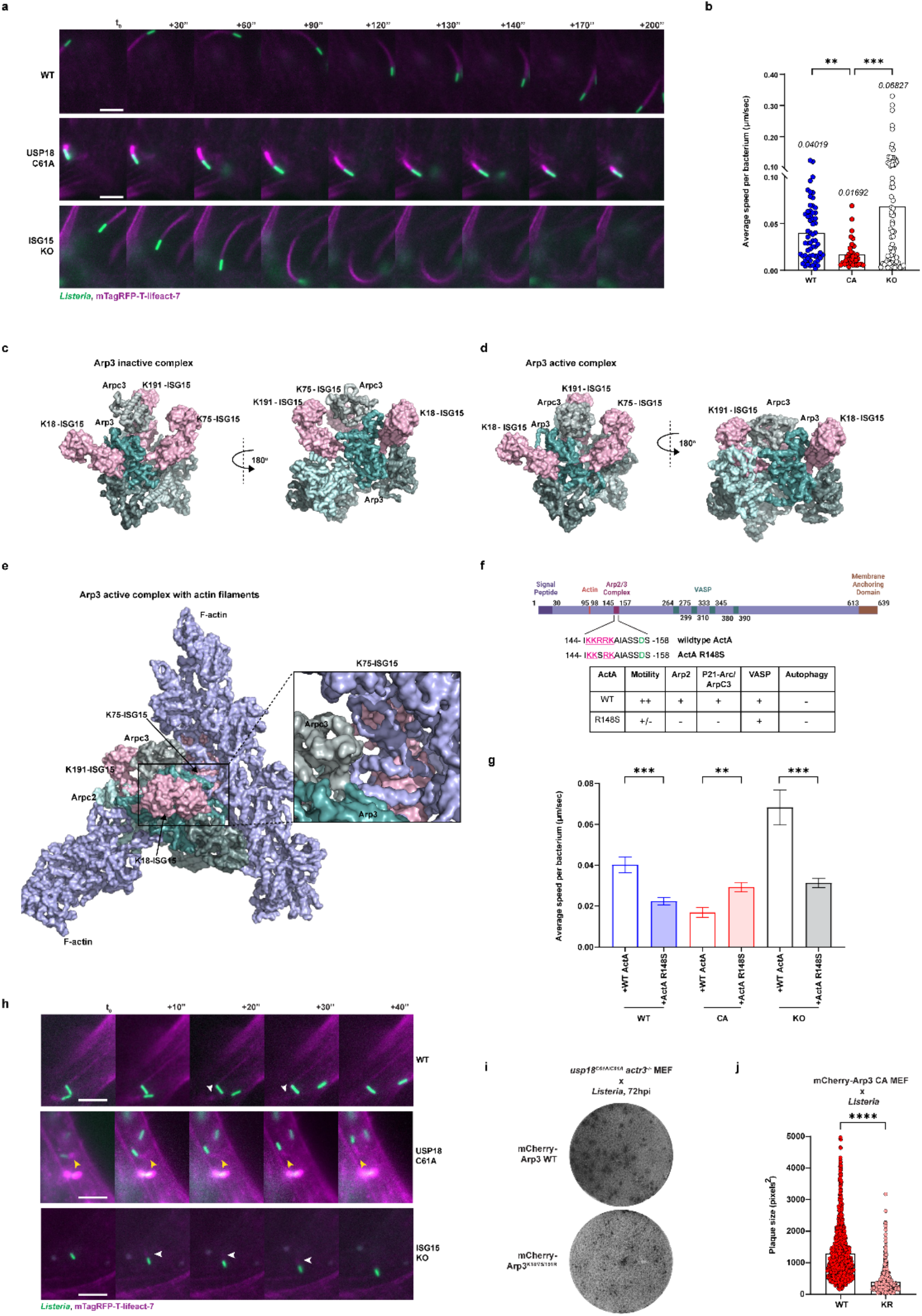
ISG15 slows Arp2/3 nucleation and bacterial comet tail speed. **a** Time-lapse microscopy of *Lm* GFP-EGD (green) infected mTagRFP-T-lifeact-7 (magenta) expressing wildtype, USP18^C61A/C61A^, or *isg15^−/−^* MEFs at 4 h post infection imaged every 10s. Scale bar, 5 μm. SI movies 1-5. **b** Quantification of average speed per bacterium tracked from 3 independent experiments (WT n=61, CA n=35, KO n=75), analyzed by Kruskal-Wallis test with Dunn’s post hoc. **c-e** Model of ISG15 (in rose) C terminus attached to Lys18, Lys75, and Lys191 of Actin-related protein 3 (Arp3) in the inactive Arp2/3 complex PDB ID: 6YW7 (**c**); the active Arp2/3 complex PDB ID: 7AQK (**d**); the active Arp2/3 complex with branched actin filaments PDB: 7AQK (**e**). **f** Schematic of ActA domains with interacting proteins; inset amino acid sequence mutated to abrogate Arp2/3 recruitment and comparison of R148S with WT for motility and autophagy; created with Biorender.com. **g** Quantification of average speed of motility per bacterium of wildtype bacteria and *Lm* EGD *DactA* complemented with ActA R148S which unstably recruits the Arp2/3 complex tracked from three independent experiments (WT n=191, CA n=203, KO n=164) by Brown-Forsythe ANOVA test with Dunnett’s T3’s post hoc. **h** Representative time-lapse images of *Lm* EGD *DactA*::ActA R148S in mTagRFP-T-lifeact-7 (magenta) expressing wildtype, USP18^C61A/C61A^, or *isg15^−/−^* MEF at 4 h post infection imaged every 10s. Scale bar, 5 μm. White arrows point to inconsistent actin tails. Yellow arrows highlight normal consistent bacterial actin tails. See also SI movies 6-8. **i** Representative *Lm* 10403s plaques in USP18^C61A/C61A^ mCherry-Arp3 and mCherry-Arp3 3KR at 3 DPI. **j** Plaque size (in square pixels) from 3 independent experiments (WT n=1073, KR n=768), analyzed with Kolmogorov-Smirnov’s t-test. In **b, g, i** and **j** asterisks indicate p values with **p < 0.01, ***p < 0.001, and ****p<0.0001. Raw data are available in the source data file.

To better understand how ISGylation could alter Arp2/3 complex function, we generated a structural model of the Arp2/3 complex with ISG15 docked on all three identified lysine modification sites (K18, K75, and K191) of Arp3 (Fig.3c-e) in the inactive, active, and actin filament bound conformations of a recent cryo-EM structure^35^. Based on the model, the observed sites of ISGylation of Arp3 could sterically hinder the interaction between the Arp2/3 complex with F-actin and may even lock Arp2/3 into an inactive conformation (Fig.3c). Thus, we hypothesize that local concentrations of modified Arp2/3 could slow actin nucleation and trap the Arp2/3 complex on the bacteria surface.

In order to assess the dependence of slowed motility in cells with enhanced ISGylation on Arp2/3 recruitment, we generated a single amino acid substitution in ActA (Extended Data Fig.3a), the bacterial virulence factor which is necessary and sufficient for *Lm* actin-based motility^36^. We mutated arginine 148 to serine (R148S), which was previously shown to disrupt stable recruitment of the Arp2/3 complex^37,38^ (Fig.3f). We reasoned that if ISG15 modification of Arp3 were responsible for restricting bacterial speed, a mutant that cannot effectively recruit the Arp2/3 complex could rescue bacterial actin-based motility in conditions of enhanced ISGylation (Fig.3g-h, SI movie 6-8). Indeed, *Lm* expressing the R148S ActA mutant displayed impaired motility in WT cells as reported. This mutant also slowed motility in ISG15-deficient cells (Fig.3g-h); however, bacterial speed was nearly restored to wildtype levels in cells with enhanced ISGylation when association between ActA and the Arp2/3 complex was perturbed (Fig.3g-h). We next sought to determine whether *Lm* EGD *DactA::actA* R148S could restrict plaque size following infection in CA monolayers. Since the mutation in ActA reduces the capacity of bacteria to spread, we allowed plaques to form for four days post infection and found that plaque size in enhanced ISGylation with this mutant is equivalent to that of wildtype cells (Extended Data Fig.3b-c) whereas plaque size remained larger in ISG15-deficient cells. Taken together, disrupting the interaction between ActA and the Arp2/3 complex rescues ISG15-slowed bacterial speed and reduces spread.

Our data suggest that ISG15 modification of the Arp2/3 complex slows actin-nucleation of pathogen comet tails. Since Arp3 has additional ISG15 sites in CA mice relative to WT in our proteomics analysis, we complemented CRISPR/Cas9 generated Arp3 deletion clones with either WT mCherry Arp3 or a non-ISGylatable mCherry Arp3 3KR with all three lysine targets mutated to arginine (K18R, K75R, and K191R). Once complemented the cells express slightly less mCherry Arp3 than endogenous Arp3 but the 3KR mutant appears to be stable with wildtype equivalent protein expression levels (Extended Data Fig.3d). We assessed cell-to-cell spread after infecting mCherry Arp3 cells with *Lm* and found that expressing a non-ISGylatable Arp3 was sufficient to reduce plaque size in cells with otherwise enhanced ISGylation (Fig.3i-j). Following Arp3 deletion and complementation, the infected cells did not detach but rather the plaques appeared as darker areas of crystal violet (Fig.3i), reminiscent of Herpes simplex virus plaques in highly adherent cell lines like HaCaT^39^. We therefore validated that the plaques were positive for the *Lm* secreted effector InlC^40^, which is frequently used as a surrogate for bacterial infection^41^. InlC staining confirms that the darker crystal violet plaques correspond to *Lm* infected cells (Extended Data Fig.3e). Taken together, enhanced ISGylation slows bacterial actin-based motility, whereas the lack of ISG15 increases speed. Importantly, a bacterial mutant which destabilizes Arp2/3 rescues WT speed and restricted spread. In addition, a non-ISGylatable mutant of Arp3 is sufficient to reduce spread pinpointing the critical role of ISG15-Arp3 in restricting pathogen spread.

### Dysregulated ISGylation of Arp2/3 complex promotes bacterial spread

Our data indicate that enhanced ISGylation slowed bacterial speed and restricted *Lm* and VacV comet tail length, by all accounts this should restrict pathogen spread. At three days post infection and in vivo, *Lm* paradoxically takes advantage of enhanced ISGylation whereas VacV cannot. Thus, we next sought to determine the molecular mechanism of enhanced spread and increased mortality in *Lm* infected CA cells and animals. In order to reconcile these disparate observations, we performed a time course of infection using fluorescence microscopy. Strikingly, at twelve hours post infection we observed groups of bacteria with bundled or shared actin tails only in CA cells (Fig.4a-b). We hypothesized that these bundles, which we termed bacterial “bazookas,” could enhance spread by bolstering force and increasing seeding in neighboring or distant cells since *Lm* cell-to-cell spread is heterogenous and dominated by pioneer bacteria^42^. Thus, we sought to understand how these unique bacterial structures form. Timelapse microscopy at eight hours post infection revealed that under conditions of enhanced ISGylation, *Lm* formed and retained actin tails on both daughter cells, whereas under WT conditions one pole typically dominates (Fig. 4f, SI movie 9-12). Furthermore, both tails generated force, resulting in bacteria that initially spiral in place rather than moving forward (Fig.4c, top panel; SI movie 9). Based on our live imaging, we also hypothesized that daughter bacteria have difficulty separating from each other following formation of actin tails in cells with enhanced ISGylation (Fig.4c); inhibition of bacterial cell division has been reported to occur with increased force on the bacterial surface from the eukaryotic cytoskeleton^43^. Notably, bacteria in the KO cells split apart much more rapidly (Fig. 4f, SI movie 12). In CA cells, we also observe that the two daughter bacteria start to move in the same direction, and some even catch other connected daughters resulting in groups of four or more bacteria with a common actin tail (Fig.4c, bottom panel; SI movie 9). We suspect that the combination of daughter cells that move together based on slow but stable actin polymerization each with a robust tail, leads to formation of bazooka bacteria that can spread farther and seed the next cell with increased individual bacteria (Fig.4c). We therefore quantified instances of daughter bacteria that were not separated but both retained actin comet tails. Up to 1% or more of bacterial cells had two tails and were still connected in CA cells whereas such bacteria were very rare in WT or KO cells (Fig.4d-e). We compared the separation time of daughter cells in all three cell lines and found that in cells with enhanced ISGylation, daughter bacteria remained attached to each other for longer (Fig. 4f; SI movie 10-12). To determine the dependence of ISG15-Arp3 on bacterial bazookas, we infected cells expressing non-ISGylatable Arp3 (3KR) and found that bacteria no longer retain two comet tails on connected daughter cells (Extended Data Fig.4b). Interestingly, however, instances of connected daughter bacteria still remained high relative to WT cells (Extended Data Fig.4a). Therefore, the lack of double Arp3 tails was sufficient to restore restricted WT spread in cells with enhanced ISGylation (Fig.3i-j). In summary, our data suggest that ISGylation of the Arp2/3 complex is an anti-pathogenic mechanism which restricts viral spread but when dysregulated can aid and abet *Lm* spread. Unlike viruses, the bacteria can divide while still attached to both comet tails forming a group which provides the potential of spreading farther and seeding more bacteria into recipient cells, resulting in enhanced mortality in mice due to unchecked spread in the liver.

**Figure 4:**
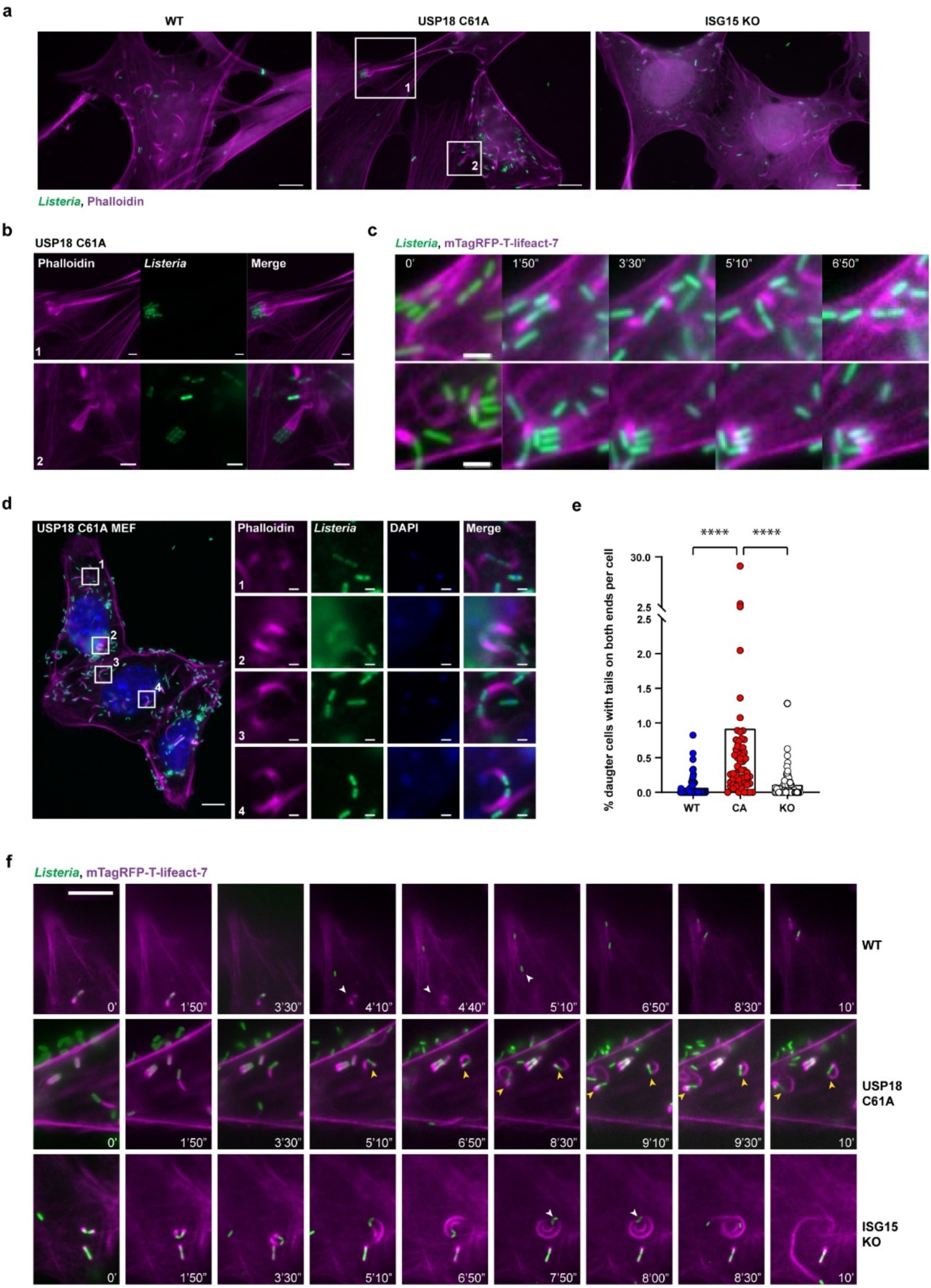
*Lm* moves into neighboring cells as a group in USP18^C61A/C61A^ cells. **a** representative image of *Lm* EGD infected wildtype, USP18^C61A/C61A^, or *isg15^−/−^* MEFs at 12 h post infection. Actin stained with phalloidin (magenta), bacteria (green); scale bar,10 μm. **b** Representative images of grouped “bazooka” bacteria with bundled actin tails formed in USP18^C61A/C61A^ MEFs at 12 h post infection; scale bars, 2 μm. **c** Time-lapse of *Lm* GFP-EGD (green) infected USP18^C61A/C61A^ MEFs expressing mTagRFP-T-lifeact-7 (magenta) at 8 h post infection, imaged every 10s; scale bar, 2 μm. See also SI movie 9. **d** Representative image of *Lm* GFP-EGD infected USP18^C61A/C61A^ MEFs at 12 HPI. Insets demonstrate bacteria with actin tails from two connected daughter cells. Actin (magenta), bacteria (green), and nuclei (blue). Scale bars in image and insets are respectively 10 μm and 1 μm. **e** The percentages of bacteria that fail to separate from each other post division with actin comet tails on both daughter cells at 12 h post infection in wildtype, USP18^C61A/C61A^, or *isg15^−/−^* MEFs; 60 individual fields from three independent experiments were counted (>459 total cells and >5624 bacteria were counted per genotype), ****p<0.0001, Kruskal-Wallis test with Dunn’s post hoc. **f** Time-lapse of *Lm* GFP-EGD (green) infected mTagRFP-T-lifeact-7 (magenta) expressing wildtype, USP18^C61A/C61A^, or *isg15^−/−^* MEFs at 8 h post infection imaged every 10s. White arrows point to bacterial division events in wildtype and *isg15^−/−^* MEFs. Yellow arrows highlight connected daughter bacteria in USP18^C61A/C61A^ MEFs that fail to separate from each other post division and retain actin comet tails on both daughter cells; scale bar, 10 μm. See also SI movies 10-12.

### ISG15 modulates cell intrinsic actin cytoskeleton dynamics

Since the Arp2/3 complex is an essential actin filament nucleator we subsequently assessed if ISGylation of the complex broadly affects cytoskeletal dynamics following infection^44^. We previously observed that our CA MEFs exhibit increased basal ISGylation which is absent in WT cells^26^. Therefore, we compared all three cell lines using STED microscopy and live imaging. Cells with enhanced ISGylation exhibited increased cortical actin thickness (Fig.5a). Whereas WT MEFs exhibit sharp, thin cortical actin more typical of fibroblasts and KO cells display diffuse cortical actin (Fig.5a). Since the cortical actin recapitulated properties of *Lm*-induced comet tails, we next sought to determine if primary mouse keratinocytes isolated from neonatal mice shared the same morphology as transformed MEFs (Fig.5b). While the cortical actin between wildtype and CA keratinocytes does not appear to be overtly different, likely due to the lack of ISG15 induction in primary skin without infection, the KO keratinocytes recapitulated the diffuse cortical actin of the fibroblasts. In terms of functional phenotypic differences, CA fibroblasts are resistant to trypsin, while the KO fibroblasts are sensitive and detach minutes earlier than wildtype (Fig.5c-d, Extended Data Fig.5a-b). We also found this to be the case with numerous ISG15-deletion or ISG15 knockdown human cancer cell lines that we have generated in the laboratory using CRISPR/Cas9 (data not shown).

**Figure 5:**
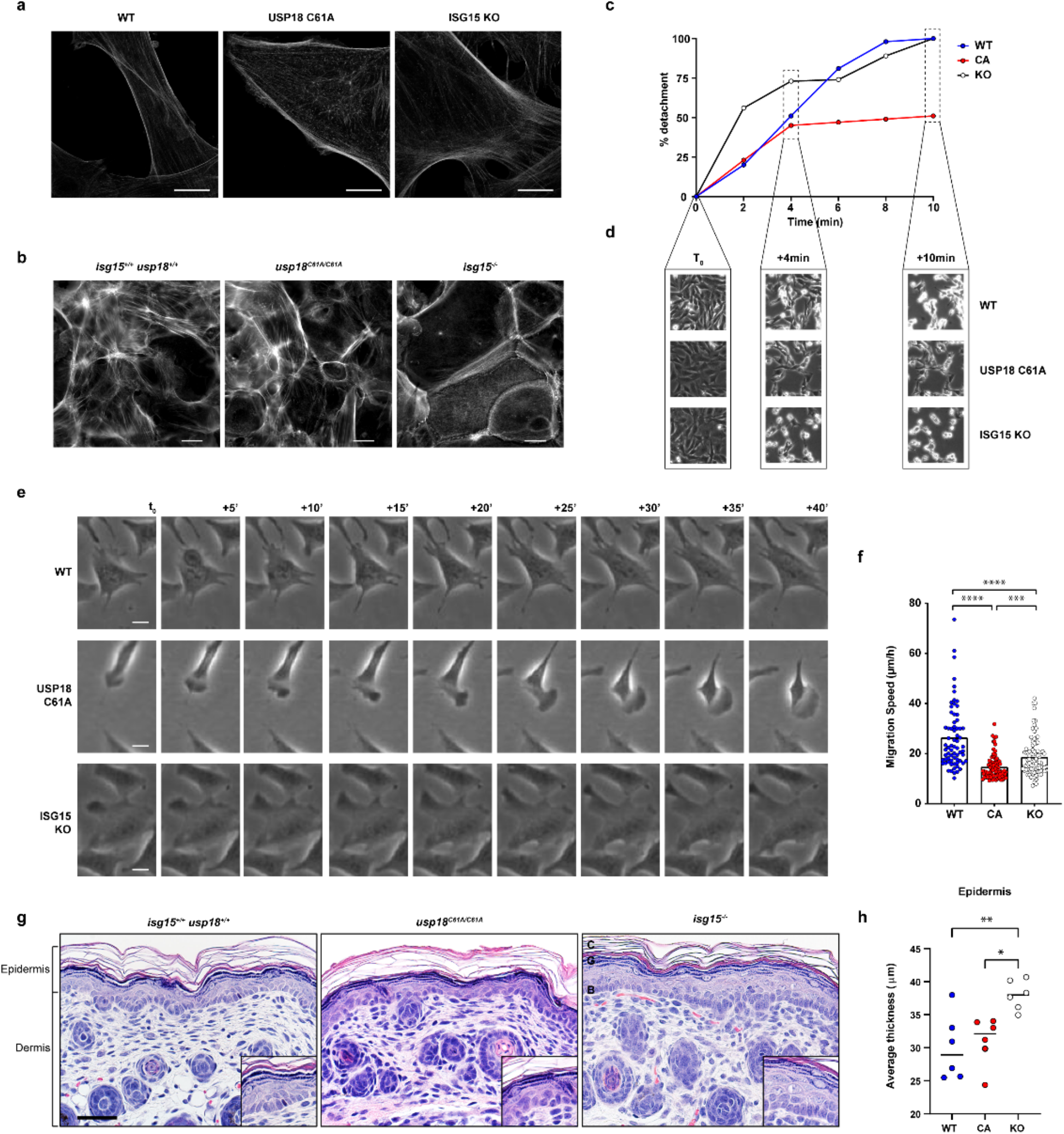
ISGylation of Arp2/3 complex affects cortical actin, adhesion, and motility in vitro and in vivo. **a** Representative STED images demonstrate differences in cortical actin thickness of wildtype, USP18^C61A/C61A^, and *isg15^−/−^* MEFs; actin (phalloidin) in white; Scale bars, 10 μm. **b** Actin in primary keratinocytes from wildtype, USP18^C61A/C61A^, or *isg15^−/−^* neonates. Actin (phalloidin) in white. Scale bars, 20 μm. **c** Quantification of average sensitivity to 1 mM EDTA/PBS in wildtype, USP18^C61A/C61A^, or *isg15^−/−^* MEF cultured on fibronectin coated surface. **d** Representative images from indicated timepoints of detachment of wildtype, USP18^C61A/C61A^, or *isg15^−/−^* MEFs. Video was sampled one frame every 20 seconds for ten minutes (6 frames/second). **e** Representative image of migration pattern of wildtype, USP18^C61A/C61A^, or *isg15^−/−^* MEF on fibronectin; scale bars, 10 μm, see movies S13-16. **f** Quantification of average migration speed of wildtype, USP18^C61A/C61A^, or *isg15^−/−^* MEF on fibronectin from six-hour videos 1 frame/min; cell positions were tracked every 6 frames; WT n=69, CA n=80, KO n=88 from two individual experiments, Kruskal-Wallis test with Dunn’s post hoc. **g** Representative histological images of neonatal skin, H and E stain, (Postnatal Day 1) from wildtype, *usp18^C61A/C61A^*, and *isg15^−/−^* mice. Strata basale (**B**), granulosum (**G**) and corneum (**C**), bar = 44 μm. **h** Quantification of epidermal thickness from six neonates from each genotype; one-way ANOVA with Tukey’s post hoc test. In **f** and **h** asterisks indicate p values with *p < 0.05, **p < 0.01, ***p < 0.001, and ****p < 0.0001. Raw data are available in the source data file.

Since the Arp2/3 complex can affect motility in a variety of cell types we quantified and compared migration speed in all three fibroblast lines. When compared to WT fibroblasts, cells with enhanced ISGylation migrated significantly (p<0.0001) more slowly (Fig.5e-f), however paradoxically the ISG15-deficient fibroblasts also migrated more slowly. In order to better understand the effect of ISG15 on cell migration we observed cells using live microscopy (Fig.5e, SI movie 13-16). CA MEFs formed long-lasting lamellipodia that resembled keratinocytes or migrating immune cells with a trailing uropod, in stark contrast to the classic filopodia-dominated motility observed in wildtype fibroblasts (Fig.5e, SI movie 13-14). The KO cells meanwhile produced large waves of unstable lamella which rapidly formed and appeared to collapse on opposite sides of the cell (Fig.5e, SI movie 15). We hypothesize that motility is affected since the cell cannot move forward due to attempts to move in two directions at once or to double back on a circuitous path. Thus, the effect we observe in actin comet tails appears to be broadly applicable to Arp2/3 cytoskeletal activity. To assess the specific role of Arp3 ISGylation in cell motility, we imaged CA Arp3-deletion cells complemented with WT mCherry Arp3 or non-ISGylatable 3KR mCherry Arp3 (Extended Data Fig.5c). The non-ISGylatable Arp3 cells no longer exhibit stable lamella like their WT Arp3 mCherry counterparts but instead they have persistent filopodia and take longer to attach (Extended Data Fig.5d-e, SI movie 17-19). Taken together, ISG15 modification of Arp3 stabilizes Arp2/3 nucleation leading to slowed motility and stable lamella, the reversal of which switched the cell periphery back to persistent fibroblast-like filopodia.

Intriguingly, recent patient case studies linked human ISG15 deficiency with ulcerating skin lesions^15,16^. These reports focused on aberrant signaling and mitochondrial metabolism in patient cells and keratinocyte cell lines but did not assess the contribution of the cytoskeleton in ISG15 deficiency. We hypothesized that ISG15 modification of actin-nucleating factors bolsters skin barrier integrity, and therefore examined the anatomy of the skin from WT, CA, and KO neonates. We reasoned that since the patients experienced lesions during early childhood that mouse neonates could best recapitulate the human patient data. Indeed, the ISG15 homozygous knock out animals have a thickened epidermal layer characterized by a proliferation or aberrant migration of large cells in the stratum granulosum (Fig.5g-h, Extended Data Fig.5g-i). While keratinocytes from ISG15-deficient mice (Fig.5b) appear larger, potentially due to effects on cell death or senescence, all of the cells displayed diffuse cortical actin, which had not previously been observed. Statistically, ISG15-deficient neonates had a significantly (p=0.0062, p=0.0164) thicker epidermis than WT and CA mice, suggesting that the absence of ISG15 can affect barrier integrity during development but that enhanced ISGylation likely requires ISG15 induction through interferon induction or infection to observe an effect on migration or epithelial integrity. Collectively, these data demonstrate that ISG15 is a key modulator of the actin cytoskeleton in metazoan development. Importantly, many of the described patient phenotypes are not conserved in mice including interferonopathy in the brain, however the absence of ISG15 modification of actin and actin modulating proteins clearly affects the actin cytoskeleton in neonatal mice, highlighting a conserved role for ISG15 that correlates with a wound healing deficiency in human patients who lack ISG15.

## Discussion

In this study, we uncovered a novel mechanism of host restriction of both viral and bacterial pathogens through modification of the Arp2/3 complex by ISG15. When properly regulated ISGylation of Arp2/3 is anti-microbial and enhanced ISGylation is protective against viral pathogens, resulting in antiviral resistance in USP18^C61A/C61A^ mice. Surprisingly, bacteria can take advantage of enhanced ISGylation by dividing while associated with stabilized actin comet tails, ultimately spreading farther into hepatocytes as groups which results in increased animal mortality relative to wildtype. By tracking *Lm* comet tail speed, we found that ISGylation slowed but stabilized actin nucleation whereas bacterial comets in cells without ISG15 rapidly nucleated actin filaments. Importantly, a single point mutation in ActA, which renders bacteria unable to stably recruit the Arp2/3 complex or a non-ISGylatable mutant of Arp3, can rescue ISGylation-induced impeded motility revealing the central role of ISG15-Arp2/3 in pathogen restriction. Finally, we uncovered changes induced by ISGylation to cell motility, morphology, and epithelial integrity and found that ISGylation slows motility and the lack of ISG15 results in altered epidermal thickening and increased sensitivity to trypsin, which correlates with defects in wound healing in ISG15-deficient patients.

### The role of free ISG15 versus ISGylation in infection

ISG15 plays distinct roles upon secretion^2,45^, non-covalent binding^3,46^, or conjugation^47^ which can counterbalance^48^, compete with each other^49^, or co-occur in the same cell or in different cell types in the same tissue. Murine models of enhanced or absent ISGylation can distinguish between separate functions however not all ISG15-associated phenotypes are conserved in mice. For instance, USP18 stabilization of ISG15 does not occur in mice^50^, thus certain aspects of the patient phenotypes could not be studied in mouse models. Here we focused on ISG15-dependent actin dynamics, which we characterized in mouse and human cells, in vivo in barrier integrity in neonates and following *Lm* infection in the liver. We found that ISG15 modification of the Arp2/3 complex affects cortical actin thickness, cell motility and sensitivity to trypsin and that this role is conserved between mouse and man. Since patients with ISG15 deficiency also lack the ability to conjugate ISG15, it is possible that ISGylation contributes to their constellation of symptoms especially in epithelial tissues. Intriguingly, recent case studies of children who present with inflammatory skin lesions have frame shift deletions in the c terminal domain of ISG15, which could potentially abrogate conjugation^10,11,51^. Pessler and colleagues also observed impaired wound healing and collective migration. When cells were reconstituted with a conjugation-deficient mutant of ISG15, deficits in collagen secretion and levels of mRNA of other cytoskeletal proteins were not rescued. Therefore, it is compelling to speculate that ISGylation of actin modulators could also contribute to the observed would healing deficits in human patients. It will be critical for future work to assess the relative contributions of the role of ISG15 modification of actin and actin binding proteins and USP18 signaling in three-dimensional tissue models with epithelial and immune cells like primary human explants or a humanized ISG15 mouse model. Such models could complement existing studies in patient-derived PBMCs^2^ or fibroblasts^9^ to dissect whether the distinct roles of ISG15 in signaling and stabilization of actin filaments synergize with increased STAT signaling to provoke aberrant wound healing.

### The double-edged sword of IFN during bacterial infections

Upon challenge of the Interferon receptor-deficient (IFNAR^-/-^) mice (or IRF3-deficient mice) with *Lm*, three independent groups made the surprising observation that without Interferon signaling the bacterial burden is significantly reduced^20–22^. At the time, bacterial replication was thought to be limited due to increased apoptosis in IFNAR^-/-^ cells^52^, thus leading to rapid clearance of bacteria by immune cells. Subsequent studies characterized the role of Type I Interferon on immune cell recruitment^53^. However, more recent work has directly implicated Type I Interferon and specific downstream ISGs in cell-to-cell spread^54^. Bacterial foci were reduced in area in IFNAR^-/-^ mice in vivo and in bone marrow derived macrophage monolayers relative to wildtype, which resulted from an impaired ability to initiate actin comet tails. Some other ISGs like IFITM3 can directly promote spread by suppressing lysosomal activity^55^. We and others have shown that ISG15 restricts bacterial replication in a cell intrinsic manner when ectopically overexpressed^23^. Conversely the lack of ISG15 can result in increased spread potentially based on increased bacterial replication per cell. Our unbiased proteomics approach and mouse model of enhanced ISGylation shed light on the critical role of direct modification of protein targets in the actin-comet tail. Actin was known to be a substrate of ISG15^28^ but modification of the Arp2/3 complex has not previously been reported. We propose that ISGylation of Arp2/3 could be partly responsible for IFNAR-dependent facilitation of cell-to-cell spread through the establishment of straighter and stronger comet tails. Indeed, in other settings, there is precedent for group motility being more efficient that single cell motility in viscous fluids^56^. For Vaccinia virus, enhanced modification limits spread, as it does for *Lm* when properly regulated, but too much modification facilitates coordinated movement of connected daughter bacteria on two stabilized comet tails ushering the troupe into neighboring cells as a pack. Interestingly, *Lm* spreads using pioneer bacteria^42^, meaning that some bacteria spread farther than others in a monolayer. We posit that this mode of invasion is favored with bacterial pairs and large ISGylation-dependent groups that we called bacterial “bazookas.” The bazookas result in deeper infiltration of hepatocytes with increased seeding of the target cell than in the wildtype liver in which bacteria are contained within immune foci. Taken together, we have identified a new molecular mechanism for ISG15 restriction of pathogen spread through modification of Arp2/3.

### ISG15 dependent effects on cell motility, cancer and wound healing

The concept of the cytoskeleton as a structural determinant of cell-autonomous host defense is emerging^57^. Type III Interferon activation increases barrier integrity^58^, which can restrict pathogen blood brain-barrier crossing. ISG15 is upregulated by both Type I and Type III Interferon^59^. By modulating the levels of ISGylation even in the absence of infection, we have shown that ISGylation directly affects actin dynamics by stabilizing lamellipodia, slowing cellular motility and increasing cortical actin thickness. ISGylation of Arp2/3 when reversed rescues cortical actin thickness, reduces lamella in the periphery favoring filopodia formation and restores restricted wildtype *Lm* spread, however cellular motility remains slow and the cells remain resistant to trypsin. We also identified alpha-actinin, cofilin and actin itself as being modified by ISG15 following infection, which could contribute to ISG15-dependent effects on motility and attachment. Early work in the cancer field implicated ISG15 as a key prognostic marker for tumorigenesis and cancer^60^. Indeed, ISG15 is upregulated in a variety of human cancer lines but intriguingly Ube1L is found within a tumor suppressor locus^61^, suggesting that some cancer cell lines have increased ISG15 that is not conjugated. In subsequent studies, ISG15 upregulation slowed wound healing and increased focal adhesions^62^, however the mechanism of action which underlies this phenomenon remained elusive. Our approach made use of CRISPR/Cas9 technology in clonal populations to recapitulate enhanced ISGylation or the lack of ISG15 from several human cell lines as well as cells derived from mouse models of enhanced ISGylation or absent ISG15. By coupling genetic control of ISGylation with mapping of ISG15 targets in the liver using mass spectrometry we were able to reveal modification sites on the surface of Arp3 which serves a molecular mechanism to slow and stabilize actin filament formation. These data mechanistically link an antiviral pathway to restriction of actin-mediated host pathogen spread, thus identifying a new function for ISG15 modification. Furthermore, our work reveals implications for conditions in which ISG15 is upregulated independent of infection like cancer. Recent studies of metastatic prostate cancer implicated ISG15 as a driver of motility^63^. It remains to be determined whether ISGylation or free ISG15 underlie this phenotype. Ultimately, given the molecular mechanism we identified, the activity of proteins that mediate conjugation or deconjugation could be tuned or leveraged using chemical inhibition to slow cell motility down in the case of metastasis, or to induce an anti-pathogenic state in the context of emerging viral infections. Taken together, ISG15 modulates cytoskeletal dynamics with clear implications for establishment of an antimicrobial state and broad potential for therapeutic intervention in difficult to treat human pathologies.

## Acknowledgements

We thank Hélène Bierne, Pascale Cossart, and Rich Roller for helpful discussions. We thank Pete Rubenstein and Mark Stamnes for critical review of the manuscript. mCherry-ARP3-N-12 and mTagRFP-T-Lifeact-7 were gifts from Michael Davidson. pUS252 was a gift from Nicholas Coleman. We thank Pascale Cossart for generously providing bacterial strains and *Lm* specific antibodies. We thank Dan Portnoy for generously providing bacterial strains. We thank Sandra Pellegrini for generously providing reagents. We used the cell sorter to obtain CRISPR/Cas9 clones at the Flow Cytometry Facility, which is a Carver College of Medicine/Holden Comprehensive Cancer Center core research facility at the University of Iowa. The facility is funded through user fees and the generous financial support of the Carver College of Medicine, Holden Comprehensive Cancer Center, and Iowa City Veteran’s Administration Medical Center. For the STED imaging the authors would like to acknowledge use of the University of Iowa Central Microscopy Research Facility, a core resource supported by the University of Iowa Vice President for Research, and the Carver College of Medicine. Y.Z. is supported by the Mechanisms of Parasitism T32AI007511-27. B.R. is supported by the Training in Mechanisms of Infectious Disease T32AI007343-33. E.U. was supported by the Predoctoral Training Program in Immunology T32AI007485-27. J.T.H is supported by AI151183, AI42767, AI114543, AI167847. D.K.M is supported by the NIH (HL163556, HL152960, DK054759, HL091842). L.R. is supported by NIGMS R35GM137961. L.R. is a Stead Family Scholar in the Carver College of Medicine.

---

**Extended Data Figure 1:**
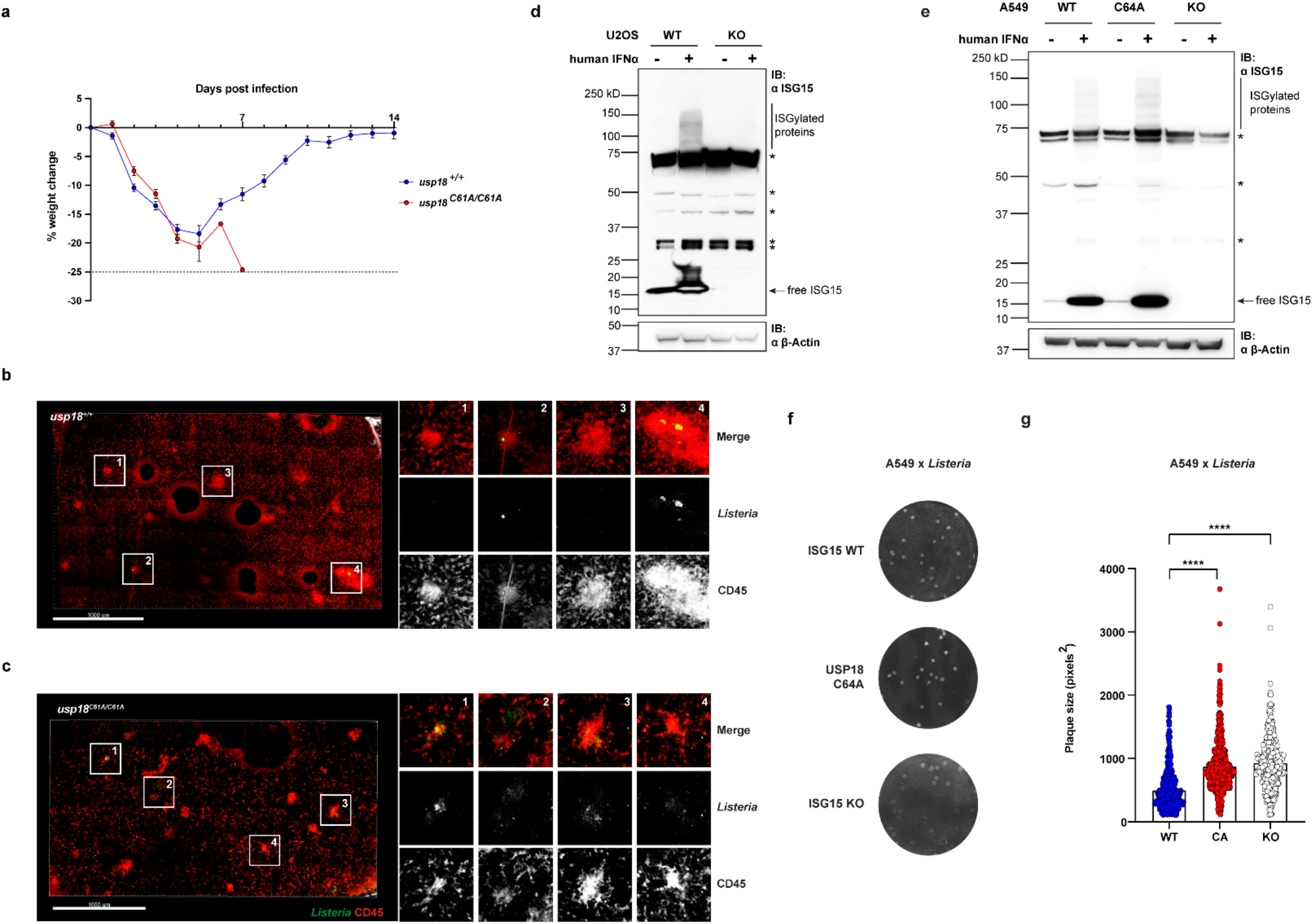
Enhanced ISGylation sensitizes mice to *L. monocytogenes* infection and promotes bacterial cell-to-cell spread. **a** Average -/+ SEM weight change in WT and *usp18^C61A/C61A^* mice infected intravenously with 1 × 10^4^ CFUs of *Lm* 10403s from 2 independent experiments (*usp18^+/+^* n = 14, *usp18^C61A/C61A^* n = 13). **b-c** Two-photon microscopy from livers of live *usp18*^+/+^ (b) and *usp18^C61A/C61A^* (c) mice infected with 5 × 10^5^ CFUs of *Lm* GFP-EGD (in green) and CD45+ cells (in green) 3 DPI. Scale bar, 1000 μm. **d** SDS-PAGE analysis of WT and *isg15^−/−^* U2OS treated with Type I IFN (1000 units/mL) for 24 h. **e** SDS-PAGE analysis of WT and *isg15^−/−^* A549 treated with IFNα (1000 U/mL) for 24 h. **f** Representative image of plaque formation of Lm EGD infection (MOI 0.0001) in WT, USP18C64A and *isg15^−/−^* A549 3 DPI. **g** Measurement of plaque size (in square pixels); data from three independent experiments (WT n=498, CA n=477, KO n=362, ****p < 0.0001), analyzed with Kruskal-Wallis test followed by Dunn’s post hoc test. In **d** and **e**, asterisks indicate background bands.

**Extended Data Figure 2:**
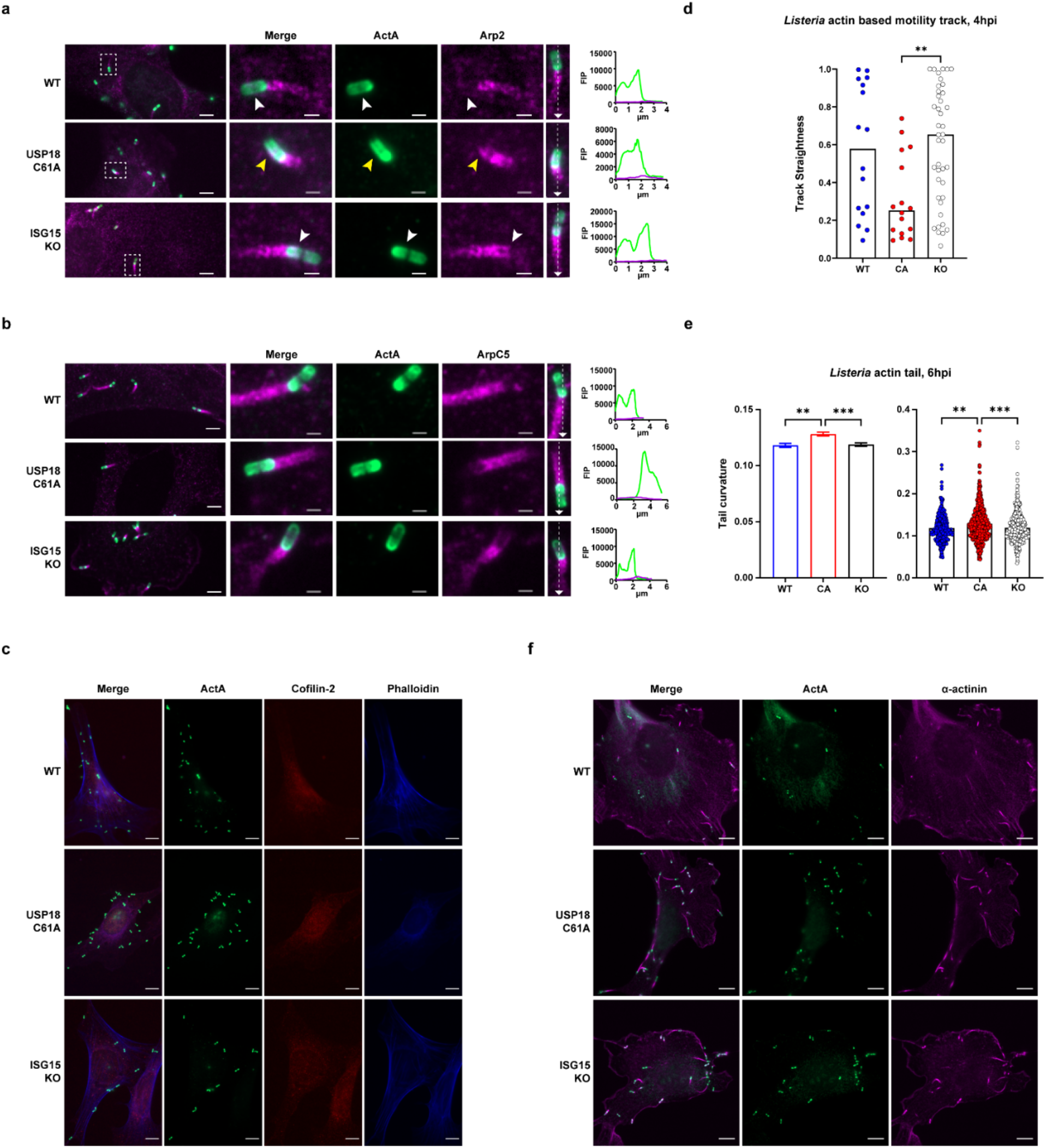
ISGylation affects recruitment of Arp2 to *Lm* bacterial surface but not ArpC5, cofilin, or alpha-actinin. **a** Arp2 comet tail in *Lm* EGD infected MEFs at 6 h post infection (MOI 10). Bacteria are shown in green, and Arp2 in magenta. Fluorescent intensity profile (FIP) was taken of the dotted line along the midline of the cell in the inset image. White arrows point to lack of colocalization. Yellow arrows highlight aberrant recruitment of Arp2 to bacterial surface. **b** Representative ArpC5 tails in *Lm* EGD infected MEFs at 6 HPI (MOI 10). Bacteria are shown in green, and ArpC5 in magenta. Fluorescent intensity profile (FIP) was taken of the dotted line along the midline of the cell in the inset image. In **a** and **b** scale bars in image and insets are 5 μm and 1 μm respectively. **c** Representative *Lm* EGD infected wildtype, USP18^C61A/C61A^, and *isg15^−/−^* MEFs at 6 HPI, a-cofilin-2 depicted in red, actin (Phalloidin) in blue, and bacteria (a-ActA) in green. Scare bars, 10 μm. **d** Analysis of actin-based motility track straightness from *Lm* GFP-EGD (green) infected mTagRFP-T-lifeact-7 (magenta) expressing wildtype, USP18^C61A/C61A^, or *isg15^−/−^* MEFs at 4 HPI; data from one representative experiment, Kruskal-Wallis test with Dunn’s post hoc (WT n=16 CA n=16 KO n=41). **e** Machine learning analysis of average curvature of actin comet tails formed by bacteria at 6 HPI in wildtype, USP18^C61A/C61A^, or *isg15^−/−^* MEFs; data represent the mean ± SEM, Kruskal-Wallis test with Dunn’s post hoc, from three independent experiments (WT n=439, CA n=661, KO n=680). **f** Representative *Lm* EGD infected wildtype, USP18^C61A/C61A^, and *isg15^−/−^* MEFs at 6 HPI, a-actinin in magenta and bacteria (a-ActA) in green. Scare bars, 10 μm. In **d** and **e** asterisks indicate p values with **p < 0.01, and ***p < 0.001. Raw data are available in the source data file.

**Extended Data Figure 3:**
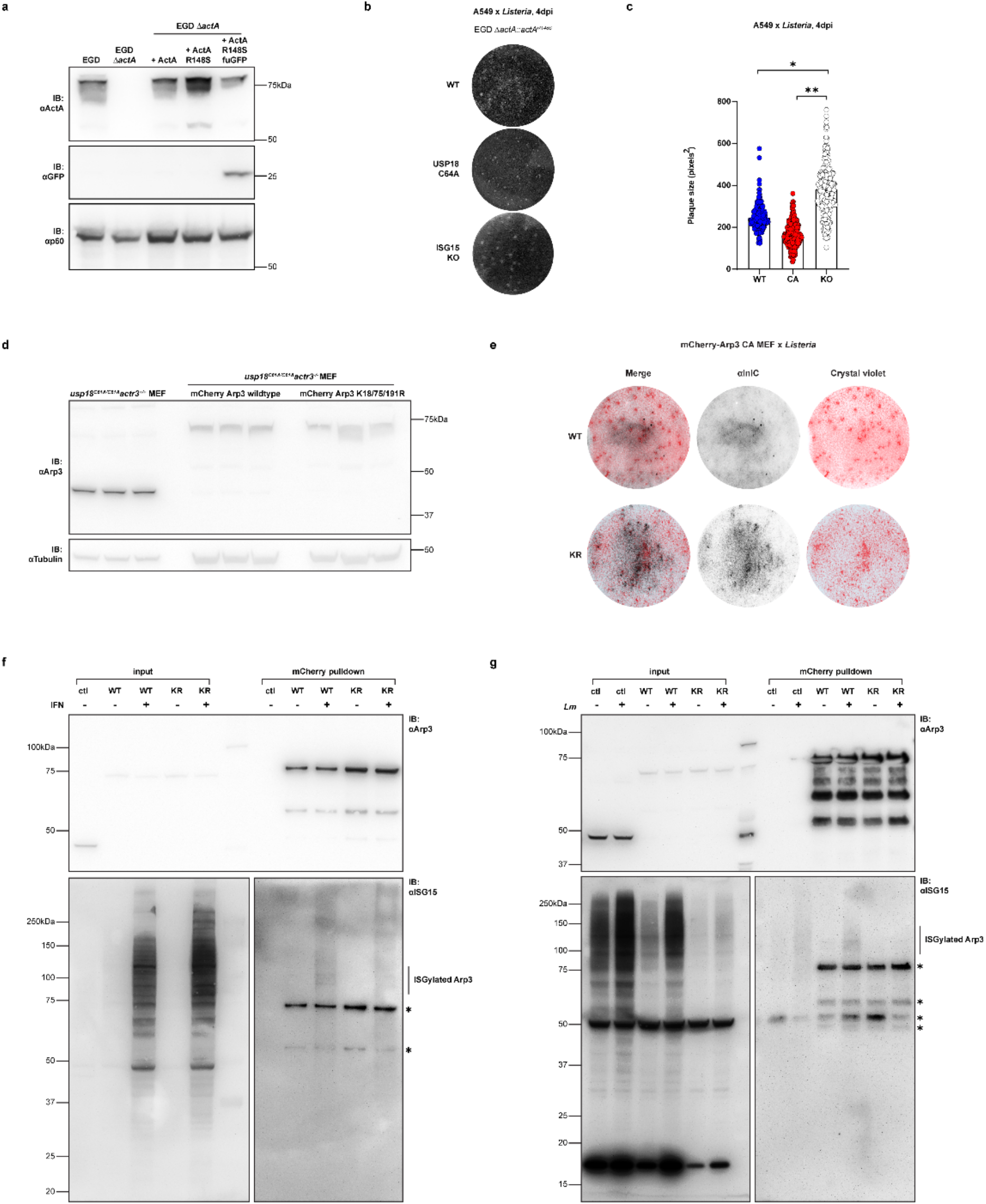
Disruption of ISG15-modification of Arp3 restricts *Lm* actin-based motility in cells. **a** Validation of ActA expression in *Lm* wild-type EGD, EGD *ΔactA*, and EGD *ΔactA* strain complemented with wildtype ActA, ActA R148S, or ActA R148S GFP constructs. **b** Representative image of plaque formation of EGD *ΔactA::actA^R148S^* infection (MOI 0.001) in WT, USP18C64A and *isg15^−/−^* A549 at 4 DPI. **c** Plaque size (in square pixels) from four independent experiments (WT n=742, CA n=945, KO n=653), analyzed with one-way ANOVA followed by Tukey’s post hoc (*p < 0.05, and **p < 0.01, raw data are available in the source data file). **d** Validation of deletion of endogenous Arp3 in usp18^C61A/C61A^ MEFs. usp18^C61A/C61A^ *actr3^−/−^* MEF clone was complemented with mCherry-Arp3 wildtype or mCherry-Arp3 that harbors lysine mutations to arginine at Lys18, Lys75, Lys191. **e** Representative image of plaque formation of *Lm*10403s in USP18^C61A^ mCherry-Arp3 and mCherry 3KR Arp3 (K18/75/191R) at 3 DPI. Infected cell monolayers were stained with a-InlC (black) and crystal violet (pink). **f** Immunoprecipitation of mCherry tag in mCherry-Arp3 complemented usp18^C61A/C61A^ *actr3^−/−^* MEFs treated with Type I IFN (1000 units/mL) for 24 h; SDS-PAGE for both Arp3 and ISG15. The experiment was repeated three times, and one representative blot is shown. **g** Immunoprecipitation of mCherry in mCherry-Arp3 complemented usp18^C61A/C61A^ *actr3^−/−^* MEFs infected with *Lm* EGD (MOI 10) for 24 h; SDS-PAGE for both Arp3 and ISG15. Blot from one representative experiment is shown. In **f** and **g**, asterisks indicate background bands from strip and reprobe.

**Extended Data Figure 4:**
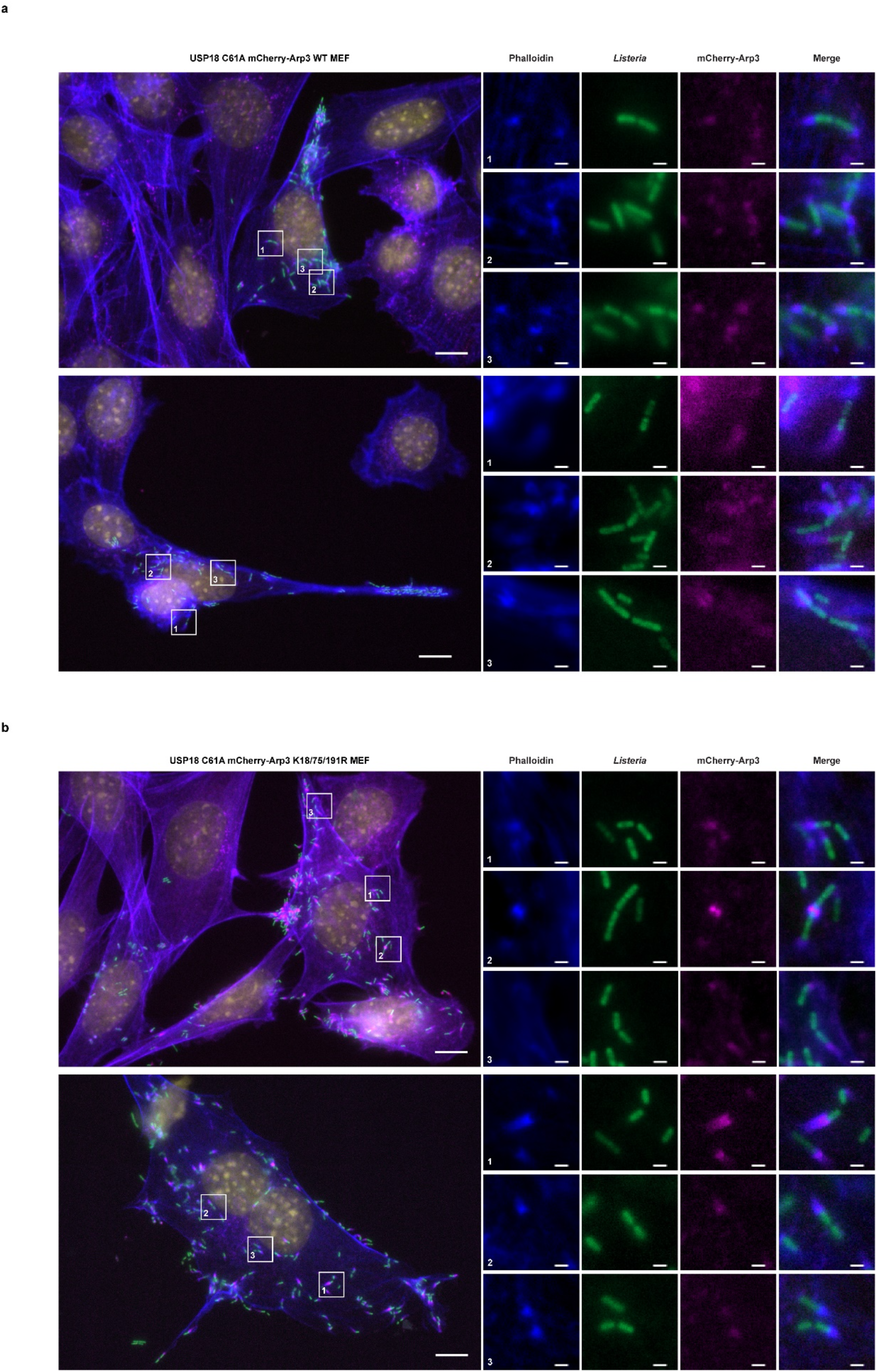
Abrogation of ISG15-modification of Arp3 disrupts stable Arp3 recruitment to both bacterial poles in cells with enhanced ISGylation. **a** Representative image of *Lm* GFP-EGD infected wildtype mCherry-Arp3 expressing *usp18^C61A/C61A^ actr3^−/−^* MEFs at 12 HPI. Insets demonstrate bacteria with actin tails from two connected daughter cells. Actin (phalloidin) in blue, bacteria (green), mCherry-Arp3 (magenta) and nuclei (yellow). **b** Representative image of *Lm* GFP-EGD infected *usp18^C61A/C61A^ actr3^−/−^* MEFs expressing mCherry-Arp3 3KR at 12 HPI. Insets demonstrate bacteria no longer associate with actin tails cells on both poles of connected daughter cells. Actin (phalloidin) in blue, bacteria (green), mCherry-Arp3 (magenta) and nuclei (yellow). Scale bars in image and insets are respectively 10 μm and 1 μm.

**Extended Data Figure 5:**
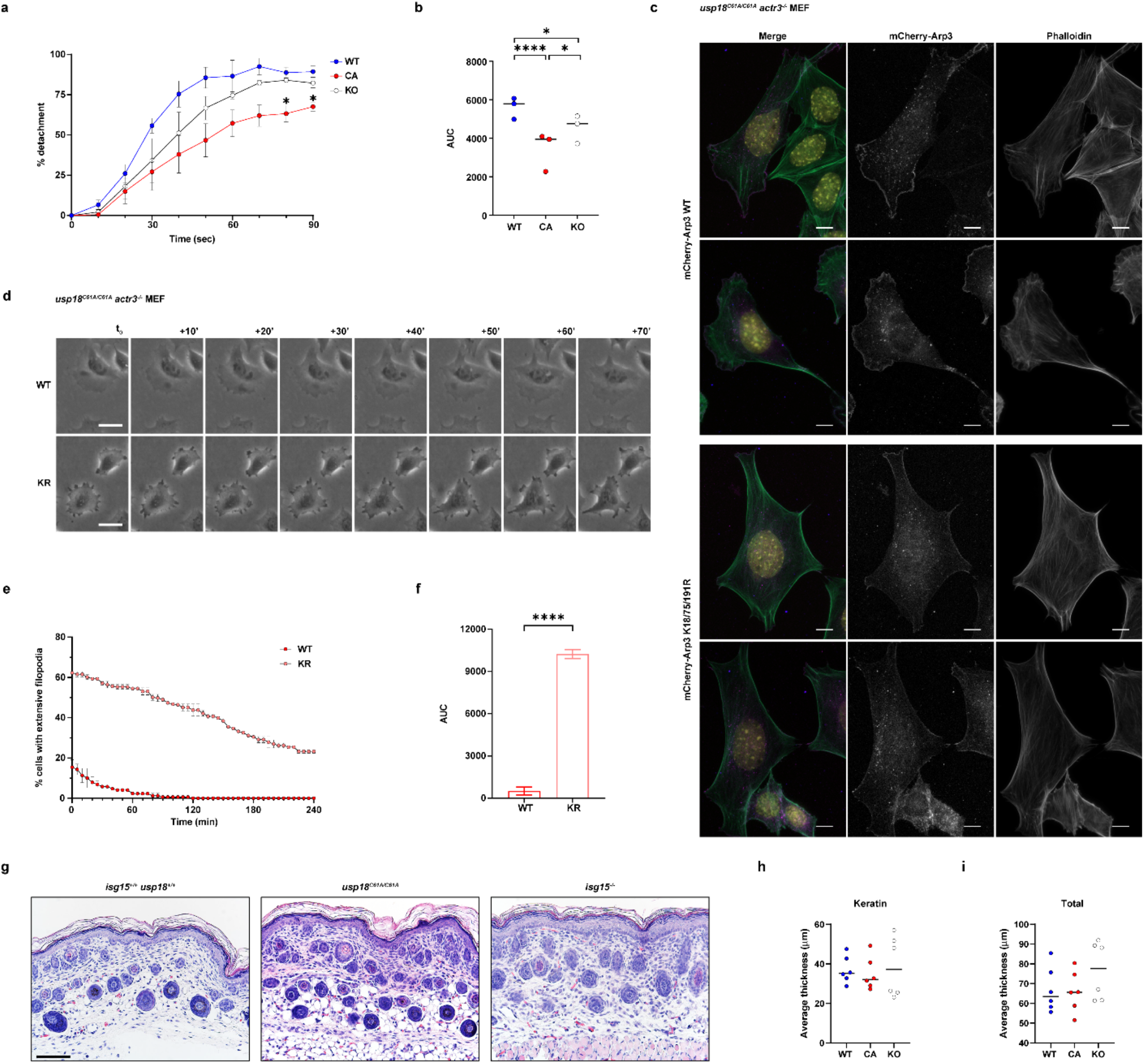
ISG15 stabilizes Arp2/3-dependent lamellipodia formation and cell adhesion. **a** Quantification of average sensitivity to 0.05% trypsin in wildtype, *Usp18^C61A/C61A^*, or *isg15^−/−^* MEFs under standard tissue culture conditions; results represent mean +/-SEM from 3 individual experiments (WT n=3, CA=3, KO=3), significance determined by Tukey’s multiple comparisons. **b** Area under the curve analysis of trypsin sensitivity in MEFs, compared with Brown-Forsythe ANOVA test with Dunnett’s T3 post hoc. **c** Representative images of actin organization and Arp3 localization in mCherry-Arp3 wildtype or Arp3 3KR expressing *usp18^C61A/C61A^ actr3^−/−^* MEFs. Actin (phalloidin) in green, mCherry-Arp3 in magenta, endogenous Arp3 (a-Arp3) in blue, and DAPI in yellow. Scale bars, 10 μm. **d** Representative image of migration pattern of Type I IFN treated (1000 U/mL for 24H) mCherry-Arp3 wildtype or Arp3 3KR expressing *usp18^C61A/C61A^ actr3^−/−^* MEFs on fibronectin; scale bars, 10 μm, see SI movies 17-19. **e** Quantification of average population of cells present with rapidly turning over filopodia-like structure (“jazz hands”). Mean -/+ SEM from 2 individual experiments, 49-95 cells were counted and tracked per genotype per repeat. **f** Area under the curve analysis of MEFs with “jazz hands”, average -/+ SEM compared with twotailed unpaired t test. **g** Representative histological images of neonatal skin (Postnatal Day 1), H and E stain, from wildtype, *usp18^C61A/C61A^*, and *isg15^−/−^* animals. Scale bar = 88 μm. **h** Quantification of keratin layer average thickness from six neonates from each genotype; oneway ANOVA with Tukey’s post hoc test. **i** Quantification of average total thickness from six neonates from each genotype; one-way ANOVA with Tukey’s post-hoc test. In **a**, **b**, and **f** asterisks indicate p values with *p < 0.05, and ****p < 0.0001. Raw data are available in the source data file.

## Methods and other online content

### Materials and Reagents

For immunoblotting, we used an anti-ISG15 antibody from Santa Cruz (F-9) at 1:200. We used an anti-GFP antibody from Invitrogen (A-11122) at 1:1000, and an anti-β-actin antibody from Sigma, Saint Louis, MO (AC-15, A5441) at 1:5000. We used an anti-Arp3 antibody from Invitrogen (PA5-17288) at 1:2000, and an anti-α-Tubulin antibody from Sigma-Aldrich (B-5-1-2, T5168) at 1:4000. We used a rabbit anti-InlC antiserum (generous gift from the Cossart Lab) at 1:1000. For immunofluorescence staining, we used an affinity purified anti-ActA antibody (P4373) at 1:100, and an anti-ActA-Nter-GST antiserum (R32) at 1:200. Both antibodies were generated in-house from rabbits (generous gift from the Cossart Lab). We used an anti-Arp3 antibody from Abcam (FMS338, ab49671) at 1:100 and an anti-Arp3 antibody from Santa Cruz (A-1, sc-48344) at 1:50. We used an anti-Arp2 antibody from Santa Cruz (E-12, sc-166103) at 1:50, an anti-ArpC5/p16-ARC antibody from Santa Cruz (C-3, sc-166760) at 1:50, an anti-Cofilin 2 antibody from Santa Cruz (D-5, sc-166958) at 1:50, and an anti-alpha actinin antibody from Sigma-Aldrich (BM-75.2, A5044) at 1:100. We used an anti-VACV 14K protein antibody generated in-house (Guerra Laboratory) and an anti-Vaccinia Virus antibody from Santa Cruz (8115, sc-58210) at 1:50. For microscopy, the aforementioned primary antibodies were revealed using Goat anti-Mouse IgG (H+L) Cross-Adsorbed ReadyProbes™ Secondary Antibody, Alexa Fluor 594 (R37121, Thermo Fisher Scientific), Rabbit IgG (H+L) Cross-Adsorbed Secondary Antibody (R37116, Thermo Fisher Scientific), and Goat anti-mouse IgG (H + L) Superclonal™ Recombinant Secondary Antibody, Alexa Fluor 647 (A28181, Thermo Fisher Scientific). Actin were labeled with phalloidin probe conjugated to either Alexa Fluor 350 dye (A22281, Invitrogen), Alexa Fluor 488 dye (A12379, Invitrogen), Alexa Fluor 594 dye (A12381, Invitrogen), or Alexa Fluor 647 dye (A22287, Invitrogen) reconstituted in methanol at 1:40. Coverslips were mounted with Fluoromount-G (Cat. 0100-01) or DAPI Fluoromount-G (Cat. 0100-20) from Southern Biotech. For super-resolution microscopy, an anti-Rabbit-IgG-Atto 647N antibody produced in goat from Sigma-Aldrich (40839) was used and actin was labeled with phalloidin probe conjugated to with Alexa Fluor 594 dye (A12381, Invitrogen). Coverslips were mounted with ProLong Glass Antifade Mountant from Invitrogen (P36982).

### Mammalian cell growth conditions

Mammalian cells were grown in DMEM with Glutamax (Gibco, Waltham, MA) supplemented with 10% fetal bovine serum (R&D Systems, Inc., Minneapolis, MN). Retrovirus was generated by transfection of 293 GP packaging cells (gift from Stipp Lab) with pBABE-puro USP18 C64A, pBABE-puro mTagRFP-T-lifeact-7, pBABEpuro-mCherry-ARP3-N-12 wildtype, or pBABEpuro-mCherry-ARP3-N-12 K all R (Plasmids listed in Supplementary Table 2) using Fugene HD (Promega, E2311). In all, 1.5 ml of collected virus was mixed with 500 μl of conditioned media of indicated cell lines supplemented with 8 μg/ml polybrene (Millipore) and applied to corresponding cells. Virus was collected and applied to cells in the presence of polybrene as described (in the protocol “Production of retroviruses using Fugene 6” from the Weinberg lab on Addgene). Cells were then selected using 2 μg/ml puromycin (Gibco, Waltham, MA) for 3 days. The population of cells that survived puromycin treatment was expanded and subsequently used for CRISPR/Cas9 deletion, knockout cell line complementation, or tested for mTagRFP-T-lifeact-7 or mCherry-Arp3 expression for timelapse microscopy. Primary keratinocytes were isolated from skin of day 1 male neonates from C57B6J mice (wildtype or USP18^C61A/C61A^ or *Isg15^-/-^*) and cultured as previously described^1^.

### Bacterial growth and strain generation

Strains used in this study are listed in Supplementary Table 1 and 2. *Listeria* strains were grown at 37°C in brain heart infusion (BHI) (Difco Laboratories, Detroit, MI). *Escherichia coli* strains were grown in lysogeny broth-Miller (LB, 1% w/v NaCl) broth at 37°C. When required, chloramphenicol was used at a final concentration of 7 μg/ml for *L. monocytogenes* and at a final concentration of 35 μg/ml for *E. coli*,kanamycin was used at a final concentration of 50 μg/ml for *E. coli*, and ampicillin was used at a final concentration of 100 μg/ml for *E. coli*.

### Cloning techniques

Primers used in this study are listed in Supplementary Table 3. Standard techniques for DNA manipulation were used. PCR was carried out using the Q5 high-fidelity PCR system (New England BioLabs Inc) according to the manufacturer’s recommendations. A GeneJET Gel Extraction and DNA Cleanup Micro kit (Thermo Scientific) and a Monarch DNA Gel Extraction kit (New England BioLabs Inc) were used for purification of DNA fragments. Restriction endonucleases and DNA-modifying enzymes were obtained from Thermo Fisher Scientific and used according to the manufacturer’s instructions. Plasmid DNA was prepared using a GeneJET Plasmid Miniprep kit (Thermo Scientific). Transformation of *E. coli* XL1-Blue (Agilent), NEB 5 alpha (New England BioLabs Inc), NEB stable (New England BioLabs Inc) and Stbl3 (Invitrogen) were accomplished by performing heat shock following the manufacturer’s instructions. Transformation of *L. monocytogenes* was accomplished by performing electroporation with a 0.1-cm cuvette using a MicroPulser apparatus (Bio-Rad) set to 25 msec, 1 pulse, and 1.8 kV as described previously^2^.

### Generation of overexpression construct

ActA integration vectors were constructed using the site-specific backbone^3^. ActA containing endogenous promoter and 5’UTR was amplified from EGD genomic DNA and R148S were introduced using primer pairs R148S FWD and REV. pAD vector was digested with EagI and SalI. Restriction fragment and amplicon were gel purified and then by Gibson assembly to generate plasmids pAD-PactA-ActA and pAD-PactA-ActA R148S. Codon optimized fuGFP sequence was provided as a gift by Nicholas Coleman and synthesized by Integrated DNA Technologies, Inc. (IDT) (Coralville, Iowa, USA). The fuGFP ActA R148S was amplified in two steps. The ActA R148S gene was amplified from pAD-PactA-ActA R148S using primers pAD-PactA-ActA FWD and REV. The fuGFP gene was amplified from gblock fragment pGroEL-fuGFP using primers fuGFP gblock FWD and REV. The two fragments were then co-ligated by SOE PCR using primers pAD-PactA-ActA FWD and fuGFP gblock REV. The final SOE PCR products, containing both ActA R148S and fuGFP under the same ActA promoter, were then cloned in pAD vector via Gibson assembly (New England BioLabs Inc). All pAD-based plasmids were verified by sequencing using primers pPL2-Rv and pPL2-Fw and were transformed into *L. monocytogenes* by electroporation. Integration into the chromosome was verified by PCR amplification using primers NC16 and PL95^2^.

Plasmid pBABEpuro-USP18 C64A was constructed as follows using the pcDNA4-UBP43 plasmid (gift from Sandra Pellegrini) as the template. C64A mutation was introduced with primer pairs USP18 C64A FWD and REV. USP18 C64A fragment was digested with SalI and gel purified, and then co-ligated with the pBABE-puro vector digested with SalI. pBABE-puro was a gift from Hartmut Land & Jay Morgenstern & Bob Weinberg (Addgene plasmid #1764; http://n2t.net/addgene:1764; RRID: Addgene_1764).^4^ Plasmid pBABEpuro-mTagRFP-T-lifeact-7 was constructed using mTagRFP-T-lifeact-7 plasmid as the template. mTagRFP-T-Lifeact-7 was a gift from Michael Davidson (Addgene plasmid #54586; http://n2t.net/addgene:54586; RRID: Addgene_54586). mTagRFP-T-lifeact-7 fragment was amplified and digested with SalI and gel purified, and then co-ligated with the pBABE-puro vector digested with SalI. Plasmid pBABEpuro-mCherry-ARP3-N-12 wildtype was constructed using mCherry-ARP3-N-12 plasmid as the template. mCherry-ARP3-N-12 was a gift from Michael Davidson (Addgene plasmid #54982; http://n2t.net/addgene:54982; RRID: Addgene_54982). mCherry-ARP3 fragment was amplified using primers mCh-ARP3 FWD and mCh-ARP3 REV, and then cloned into pBABE-puro vector, linearized with BamHI and SalI, via Gibson assembly (New England BioLabs Inc). pBABEpuro-mCherry-ARP3-N-12 K all R plasmid were generated in two steps. gblock fragment ARP3 K all R was digested with NheI and BamHI, column purified, and then co-ligated with the mCherry-ARP3-N-12 vector linearized with NheI and BamHI. mCherry-ARP3 K all R fragment was amplified with primers mCh-ARP3 FWD and mCh-ARP3 REV and then cloned into pBABE-puro vector, linearized with BamHI and SalI, via Gibson assembly (New England BioLabs Inc). All pBABEpuro-based plasmids were verified by sequencing using primers 5’ sq pBABE and 3’ sq pBABE.

### Development of CRISPR/Cas9 cell lines

Guide RNA sequences used in this study are listed in Supplementary Table 3. The ISG15 knockout cell lines were generated by using a CRISPR/Cas9 approach. Target sequences were designed with Benchling (benchling.com). Oligonucleotides were synthesized by IDT (Coralville, IA, USA) and cloned into the pX330-U6-Chimeric_BB-CBh-hSpCas9 or pSpCas9(BB)-2A-GFP plasmid. pX330-U6-Chimeric_BB-CBh-hSpCas9 was a gift from Feng Zhang (Addgene plasmid #42230; http://n2t.net/addgene:42230; RRID: Addgene_42230)^5^. pSpCas9(BB)-2A-GFP (PX458) was a gift from Feng Zhang (Addgene plasmid #48138; http://n2t.net/addgene:48138; RRID: Addgene_48138)^6^. Cells were co-transfected with the two ISG15-targeting plasmids with FuGene HD. 48 h post transfection, GFP positive cells were sorted and plated in 96-well plate for single-clone selection with Becton Dickinson FACS Aria Fusion (BD Science). ISG15 deficiency was tested by IFN-α stimulation and subsequent western blotting for ISG15.The absence of ISG15 protein expression was confirmed by immunoblotting as well as genomic PCR and next-generation sequencing of the amplicon (amplification primers for the target region are listed in Supplementary Table 3).

The USP18 C64A A549 cell line was generated by a two-step approach. First, USP18 C64A under the LTR promoter of pBABE-puro vector was expressed in wildtype A549 cells as described above. Then using a CRISPR/Cas9 approach, single-guide RNAs (sgRNAs) targeting USP18 endogenous promoter regions on the chromosome were designed using Benchling. Oligonucleotides were synthesized by IDT and cloned into the pSpCas9(BB)-2A-GFP plasmid. Cells were co-transfected with the two USP18 promoter-targeting plasmids with FuGene HD 48 h post transfection, GFP positive cells were sorted and plated in 96-well plate for single-clone selection with Becton Dickinson FACS Aria Fusion. The absence of endogenous USP18 promoter was confirmed genomic PCR and next generation sequencing of the amplicon (amplification primers for the target region are listed in Supplementary Table 3). Abolishment of enzymatic function of USP18 and increased ISGylation were tested by IFN-α stimulation and subsequent immunoblotting against ISG15.

The Arp3 knockout cell lines were generated also via a CRISPR/Cas9 approach. Target sequences were designed with Benchling (benchling.com). Oligonucleotides were synthesized by IDT (Coralville, IA, USA) and cloned into the pSpCas9(BB)-2A-GFP plasmid. Cells were co-transfected with a Arp3-targeting plasmid with FuGene HD. 48 h post transfection, GFP positive cells were sorted and plated in 96-well plate for single-clone selection with Becton Dickinson FACS Aria Fusion (BD Science). We observed that Arp3 deletion dramatically affected cell attachment and slowed down cell replication. Similar growth and attachment defect had also been reported and described by previous literatures. Thus, we immediately complimented viable candidate clones with mCherry-Arp3 constructs. Each candidate clone was split into two wells and transformed with either pBABEpuro-mCherry-ARP3-N-12 wildtype or pBABEpuro-mCherry-ARP3-N-12 K all R viruses. Arp3 deficiency was tested by western blotting for Arp3. The absence of endogenous Arp3 protein expression was confirmed by immunoblotting as well as genomic PCR and next-generation sequencing of the amplicon (amplification primers for the target region are listed in Supplementary Table 3).

### Plaque Formation of *Listeria monocytogenes* in MEFs and A549s

In all cases, cells were plated so that they were confluent on the day of the infection. *Listeria monocytogenes* strains were grown in BHI or BHI supplemented with chloramphenicol overnight at 37°C with shaking. Prior to infection, overnight cultures of bacteria were diluted in new media and grown to exponential phase (OD 0.8–1) for EGD based strains and stationary phase (OD >1.2) for strain 10403s. Then the bacteria were washed three times and resuspended in serum-free mammalian cell culture media at the indicated MOI. A fixed volume was then added to each well. Cells were centrifuged for 1 min at 201 × g to synchronize infection. The cells were then incubated with the bacteria for 1 h at 37 °C, 5% CO2. Following this incubation, the cells were washed at room temperature with 1× PBS (Gibco, 14190-144) and overlaid with medium containing 1% agarose, low melt (IBI Scientific, Dubuque, IA) and 20 μg of gentamicin sulfate per ml (a stock of 2 × medium was prewarmed at 37°C and mixed with 56°C 2% agarose immediately before use). After 3 days, plaques were fixed with 4% formaldehyde solutions (Sigma-Aldrich, #252549) in PBS at RT overnight and stained with 0.5% crystal violet solution (0.5% w/v crystal violet, 20% v/v methanol) (Sigma-Aldrich) for 1 h at RT. Plaques were visualized as clear zones in a lawn (or more intensely stained clusters in *actr3^/-^* cells) of purple cells and scanned by an EPSON perfection V500 Photo scanner. Size of the plaques were measured in ImageJ using ViralPlaque plugin^7^.

For immunostaining of plaques in USP18^C61A/C61A^ *actr3^/-^* × mCherry-Arp3 wildtype or mCherry-Arp3^K18/75/191R^ MEFs, after 3 days, plaques with 4% formaldehyde solutions (Sigma-Aldrich, #252549) in PBS at 4°C overnight, permeabilized with 0.5% Triton in PBS at RT for 15 min, and blocked with 5% BSA in 0.05% Tween-80 in PBS for 1 hr at RT. We immuno-stained *L. monocytogenes* infected cells with anti-InlC antiserum for 1 hr followed by 1 hr incubation with peroxidase-labeled anti-rabbit antibodies at 1:1000 (Kindle Biosciences, LLC). Solution of 5% bovine serum albumin (BSA, VWR) and 0.05% Tween-80 in PBS was used for the preparation of working dilutions of immuno-reagents. Cells were washed after the primary and secondary antibodies by incubating them three times for 5 min with 0.05% Tween-80 in PBS. As peroxidase substrates, we employed Kwik Quant ultra-digital ECL substrate solution (Kindle Biosciences, LLC), and imaged with Kwik Quant Imager system (Kindle Biosciences, LLC).

### Extraction of bacteria from infected cell lines

*Listeria monocytogenes* strain GFP-EGD was grown in BHI. Prior to infection, overnight cultures of bacterial strains were back diluted and grown to exponential phase (OD 0.8–1), washed three times and resuspended in serum-free mammalian cell culture media at MOI of 10. A fixed volume was then added to each T175 flask of MEFs. Cells were centrifuged for 1 min at 201 × g to synchronize infection. The cells were then incubated with the bacteria for 1 h at 37 °C, 5% CO2. Following this incubation, the cells were washed at room temperature with 1× PBS, and cell growth medium with 10% serum was added with 20 μg/ml gentamicin to kill extracellular bacteria. At 24 h post infection, cells were washed twice with 1x PBS and scraped up into 5 ml PBS per flask. Cells were gently pelleted at 800 × g for 5 min and resuspended in 4 ml 1mM NaHPO4/NaH2PO4 buffer, pH 7.0. Cells were broken with three strokes of the tight pestle, 1 ml of 1.25 M sucrose (final concentration will be 0.25 M) was immediately added to stabilize the nuclei, and then mixed with two more strokes of the pestle. The homogenates were spun at 3990 × g at 4°C for 5 min to pellet out Nuclei. The cytosolic supernatants were spun again at 8000 × g at 4°C for 3 min to obtain the bacterial/mitochondrial pellets. 1 ml 0.1% Triton in H_2_O was added to the pellets and mixed via pipetting to burst mitochondria. The final bacterial pellets were collected via spinning the triton lysates at 8000 × g at 4 °C for 3 min. The bacterial pellets were resuspended in PBS, normalized across the genotypes according to the optical density at 600nm read by an Eppendorf biospectrometer, and lysed with 2× Laёmmli buffer containing 62.5 mM Tris-HCl pH 6.8, 2% SDS, 10% glycerol, 0,002% Bromophenol blue supplemented with 20 mM DTT and 10% 2-mercaptoethonal (VWR) at 95°C for 15min for SDS-PAGE analysis.

### In vivo infection

All animal studies and procedures were approved by the University of Iowa Animal Care and Use Committee, under U.S. Public Health Service assurance, Office of Laboratory Animal Welfare guidelines. For survival studies, female C57BL/6 mice (wildtype or USP18^C61A/C61A^ or *Isg15^-/-^*) were infected intravenously via tail veil injection between 8 and 10 weeks of age with wild-type *L. monocytogenes* strain 10403S at 1 × 10^5^ CFUs per animal. Mice were assessed daily 14 days after inoculation and euthanized if they lost more than 25% of their body weight, became moribund, or dropped below a body condition score equal or less than two. For two-photon microscopy, female C57BL/6 mice (wildtype or USP18^C61A/C61A^) were infected intravenously via tail veil between 8 and 12 weeks of age with GFP-EGD at 5 × 10^5^ CFUs per animal. Mice were imaged 72 h following infection (see intravital microscopy section).

### Intravital microscopy

All images were acquired on a SP8 NLO Microscope (Leica) using a 25x / 0.95 NA water immersion objective with coverslip correction as previously described^8^. High resolution stacks of 20-35 xy sections sampled with 1-3μ z spacing were acquired at an acquisition rate of 30-60 seconds per stack and merged with Leica software. Where indicated, mice were injected with F4/80-APC (10μg, i.v.) 1 hour prior to imaging or with CD45-PE (10μg, i.v.), or Evans Blue (0.1% solution in PBS, 200μl i.v.) for vascular label immediately prior to imaging.

All animal studies and procedures were approved by the University of Iowa Animal Care and Use Committee, under U.S. Public Health Service assurance, Office of Laboratory Animal Welfare guidelines. Mice were anesthetized with Ketamine/Xylazine (87.5/12.5 mg/kg) prior to their surgical preparation for imaging. Mice were placed in dorsal recumbency on a heating pad maintained at 37°C, an abdominal anterioposterior incision was made in the skin posterior to the diaphragm. Lateral incisions were made on the left side through the skin at either ends of the earlier incision and skin reflected to the side. Subsequently, the peritoneal membrane was incised and reflected as above, to expose the liver. Mice were positioned on dorsal recumbency on the microscope base in a continuously heated (37°C) enclosed chamber (Leica). A custom tissue-suction apparatus (VueBio) was placed on the liver with 20-25mm Hg of negative pressure to gently immobilize the liver tissue against a fixed coverslip. All Images were acquired using the following Ex/Em parameters for the various proteins or fluorophores: PE 561/567-623, GFP 940/490-510, APC 1250, collagen matrix Secondary Harmonic Generation (SHG) (535-545nm). Sequences of acquired image stacks were transformed into volume-rendered, isosurfaced, 3D images and 4D time-lapse movies with Imaris Version 8.1 (Bitplane).

### Histopathology and morphometry

Skin sections were collected along the dorsum in a transverse plane. Tissues were fixed in 10% neutral buffered formalin (~1 week) and routinely processed, embedded, sectioned (~4 μm) and stained with hematoxylin and eosin (HE) stain. Tissues were evaluated by a boarded veterinary pathologist using the post-examination method of masking to group assignment^9^. High resolution digital images of dorsal skin samples were collected for each mouse and measurements (an average of 3 different samples) taken for the epidermis (stratum basale through stratum granulosum), keratin layer (stratum corneum) and total thickness (stratum basale through stratum corneum). Measurements were consistently made in regions with an intact epidermis that lacked sectioning artifacts. Samples were analyzed by one way ANOVA with Tukey’s post hoc test with significance defined at p < 0.05.

### SDS-PAGE

For the SDS-PAGE analysis of mammalian cells, cells lysed in 1% Triton lysis buffer supplemented with Pierce protease inhibitor mini tablet (Thermo Scientific, A32953). Gels were transferred using an iBLOT transfer system (Invitrogen), blocked in 5% milk for 1 h at RT, incubated with primary antibody overnight at 4 °C, washed with 0.05% Tween in PBS three times (each wash for 7 min), and incubated for 1 h at room temperature with secondary antibody coupled to HRP (Kindle Biosciences, LLC). Blots were washed again three times with 0.05% Tween in PBS, revealed using Kwik Quant ultra-digital ECL substrate solution and captured with Kwik Quant Imager system (Kindle Biosciences, LLC).

### Immunoprecipitation

Cells were lysed and protein concentration was determined with either a Bradford or BCA assay. In total, 2 mg of total protein were used for mCherry-immunoprecipitation, using RFP-Trap^®^ (ChromoTek) according to the manufacturer’s instructions. Mouse embryonic fibroblasts were treated with recombinant mouse type I interferon IFN-alpha A protein (R&D system) at 1000 U/ml or untreated for 24h or infected with *Lm* strain GFP-EGD at MOI of 10 or uninfected for 24h. At 24 h, posttreatment cells were lysed in 1x RIPA lysis buffer supplemented with Pierce protease inhibitor mini tablet (Thermo Scientific, A32953) and phosphatase inhibitor cocktails. The lysates were further treated with universal nuclease for 30 min at 4 °C (Pierce). Each sample was incubated with 25 μl of RFP-Trap^®^ (ChromoTek) bead slurry at 4 °C overnight. Beads were pulled down with a magnet (Dyna Mag2, Invitrogen) and washed five times with following the manufacturer’s instructions. Proteins were eluted from beads with 2× Laëmmli buffer containing 62.5 mM Tris-HCl pH 6.8, 2% SDS, 10% glycerol, 0,002% Bromophenol blue supplemented with 10% 2-mercaptoethonal (VWR) at 95°C for 5min for SDS-PAGE analysis.

### Fixed widefield and confocal microscopy

For *Listeria* infection, cells were plated on coverslips the day prior to an experiment. Cells were fixed in 4% PFA (Electron Microscopy Sciences, Hatfield, Pennsylvania) in PBS for 20 min at room temperature, and permeabilized with 0.5% Triton in PBS for 10 min at room temperature. Coverslips were incubated with primary antibodies followed by Alexa conjugated secondary antibodies in blocking solution for at least 1 h at room temperature. Coverslips were then mounted either in Fluoromount-G mounting medium with or without 4’,6-diamidino-2-phenylindole (DAPI, SouthernBiotech) for wide-field fluorescent imaging. Images were acquired using an inverted wide-field fluorescence microscope (Axio Observer 7, Carl Zeiss Microscopy, Germany) equipped with an Axiocam 506 mono camera and the software ZEN 2.3 Pro.

For vaccinia virus infection, cells were seed on coverslips prior to infection. Cells were infected with the vaccinia virus strain WR at MOI 0.5 for 8 h or MOI 1 for 24 h. For 8 HPI, cells were fixed in 4% PFA (Electron Microscopy Sciences, Hatfield, Pennsylvania) in PBS for 20 min at room temperature, and permeabilized with 0.5% Triton in PBS for 10 min at room temperature. Coverslips were incubated with primary antibodies followed by Alexa conjugated secondary antibodies in blocking solution for at least 1 h at room temperature. Coverslips were then mounted either in Fluoromount-G mounting medium with or without 4’,6-diamidino-2-phenylindole (DAPI, SouthernBiotech) for wide-field fluorescent imaging. Images were acquired using an inverted wide-field fluorescence microscope (Axio Observer 7, Carl Zeiss Microscopy, Germany) equipped with an Axiocam 506 mono camera and the software ZEN 2.3 Pro. For 24 HPI, cells were washed with Cytoskeleton Buffer (10mM MES, 10mM NaCl, 5mM EGTA, 5mM MgCl2, 5mM Glucose) (CB) and fixed with 3% paraformaldehyde (PFA) in Cytoskeleton Buffer. Cells were washed twice with Cytoskeleton Buffer and stored at 4°C for further processing. Cells were permeabilized with 0.25% Triton and blocked with 10% FCS. Specific monoclonal antibody for the VACV 14K protein was used and Alexa Fluor 488-conjugated mouse IgG (Invitrogen) was used as secondary antibody. Cell nuclei were stained with 4’,6-diamidino-2-phenylindole (DAPI) (Sigma). F-actin was stained with rhodamine-conjugated phalloidin (Molecular Probes). Confocal microscopy was performed using a Leica SP8 laser scanning microscope, and images were collected and processed with LAS AF software (Leica, Wetzlar, Germany).

### Stimulated Emission Depletion Microscopy (STED)

Fibroblasts were attached to #1.5 High Precision Glass Cover (Bioscience Tools). Cells were fixed in 4% PFA (Electron Microscopy Sciences, Hatfield, Pennsylvania) in PBS for 20 min at room temperature, and permeabilized with 0.1% Triton in PBS for 10 min at room temperature. Coverslips were blocked with 5% BSA supplemented with 1% goat serum in PBS at 4°C overnight.

For infected cells, coverslips were then stained with rabbit a-ActA (affinity purified clone P4373, gift from Cossart lab) for 1 h at RT. Coverslips were washed with 5% BSA in PBS for three times (each wash for 5 min) and incubated for 1 h at RT with AttoN647 conjugated anti-rabbit secondary antibodies and Alexa Fluor 594 conjugated Phalloidin. For non-infected cells, coverslips were incubated for 1 h at RT with Alexa Fluor 594 conjugated Phalloidin. Coverslips were washed again three times with 5% BSA in PBS, and subsequently mounted with Prolong Glass Antifade medium. Z-stack images were acquired using a Leica SP8 STED Super Resolution Microscope and LAS × software (Leica Microsystems, Buffalo Grove, IL).

### Time-lapse fluorescence microscopy

mTagRFP-T-lifeact-7 MEFs were seeded in 35 mm #1.5H glass bottomed dishes (ibidi, μ-Dish 35 mm, high Glass Bottom) using 1 × 10^5^ cells in media without phenol red (Gibco, 21063029). Overnight cultures of *L. monocytogenes* strains expressing GFP were diluted in Brain Heart Infusion media (BD) and grown to exponential phase (OD 0.8–1), washed three times in serum-free DMEM (Life Technologies), and resuspended in serum-free DMEM at MOI of 10. A fixed volume was then added to each well. Cells were centrifuged for 1 min at 201 × g to synchronize infection. The cells were then incubated with the bacteria for 1 h at 37 °C, 5% CO_2_. Following this incubation, the cells were washed at room temperature with 1 × DPBS, and then fed with full growth medium (with 10% fetal bovine serum) supplemented with 20 μg/ml gentamicin to kill extracellular bacteria. At 4 h or 8 h post infection, infected cells were washed 3 times with PBS and imaging was performed in media without phenol red containing a 1:100 dilution of ProLong Antifade Reagents for Live Cells (Invitrogen).

Time-lapse microscopy experiments were performed using an inverted wide-field fluorescence microscope (Axio Observer 7, Carl Zeiss Microscopy, Germany) equipped with an Axiocam 506 mono camera and the software ZEN 2.3 Pro, in an incubation chamber at 37°C with 5% CO2. For each XY position, images were acquired every 10s for 10 minutes with a 63× plan apo objective. All supplemental videos were corrected for bleaching using the stack correction function (third dimension:T, corrected each channel individually, background+decay+flicker) in Zen 2.3 Pro.

### Cell migration assay

On the day of experiment, MEFs were rinsed once and lifted with 1mM EDTA in PBS at RT (~3-5mins for wildtype or *isg15^-/-^* MEF, and ~10min for *usp18^C61A/C61A^* MEF). After centrifugation, cells were resuspended in full media and seeded at 1.2 × 10^5^ cells to 35mm glass-bottomed cover dishes coated with 20 μg/ml fibronectin (Corning, #354008). Cells were allowed to attach for 2h, cultures were imaged on a Leica DMIRE2 inverted microscope (Deerfield, IL) in a stage incubator (20/20 Technology, Wilmington, NC) providing a humidified 5% CO2, 37 °C atmosphere. For USP18^C61A/C61A^*actr3^-/-^* MEFs, cells were treated with mouse IFNα overnight, lifted with 0.05% trypsin/EDTA at RT and seeded at 2.0 × 10^5^ cells. Cells were allowed to attach for 1.5h prior to imaging. OpenLab software (Improvision, Lexington, MA) running on an Apple iMac computer-controlled illumination and image acquisition (Cupertino). Images were acquired at a rate of 1 frame/min for 6 h using a Hamamatsu ORCA-285 CCD camera (Bridgewater, NJ) and a 20 C Plan phase objective. ImageJ software was used to record the XY position of cell centroids in the frames 6 min apart, using the mTrack plugin^10^. Every cell in each field was followed for as long as it remained in view. All cells that could be tracked for at least 1 h were included in the final calculation of velocity (typically 40–60 cells per experiment).

### Cell detachment assay

Cells were grown on 35mm glass-bottomed cover dishes to 90-100% confluent at the day of experiment. Cells were imaged and observed for 10min after flush of 1mM EDTA/PBS on a Leica DMIRE2 inverted microscope (Deerfield, IL) in a stage incubator (20/20 Technology, Wilmington, NC) providing a humidified 5% CO2, 37 °C atmosphere. OpenLab software (Improvision, Lexington, MA) running on an Apple iMac computer-controlled illumination and image acquisition (Cupertino). Images were acquired at a 20s-sampling-interval for 20 min using a Hamamatsu ORCA-285 CCD camera (Bridgewater, NJ) and a 20 C Plan phase objective. Cells were imaged and observed for 5min15s after flush of 0.05% trypsin/EDTA using an inverted wide-field fluorescence microscope (Axio Observer 7, Carl Zeiss Microscopy, Germany) equipped with an Axiocam 506 mono camera and the software ZEN 2.3 Pro, in an incubation chamber at 37°C with 5% CO2). Images were acquired at a 5s-sampling-interval for 5 min 30s. CellProfiler software^11^ was used to record the total surface area occupied by cells at each time point, with the pixel based identification pipeline^12^. The changes in total occupied surface area were tracked and recorded in ImageJ. In short, images were auto-threshold, and surface area were extrapolated with particle measurements (size limited to over 50 pixels^2^).

### LC-MS/MS data analysis

ISG15 modified sites were identified as previously described^13^. Of the 929 ISG15 modified sites identified, 13 sites identified in the actin-comet tail proteome^14^ were sorted based on the ANOVA intensities previously described (Figure 2a) and shown in a heatmap (Fig.1f) after non-supervised hierarchical clustering. Column and row clustering were determined based on imputed values from a normal distribution.

### Modeling modified Arp3 in the Arp2/3 complex

To model ISGylated Arp3 in the Arp2/3 complex, first we used the program Coot to add the missing residues, RGG, to the C-terminus of the ISG15 PDB file, 1Z2M. Pymol was then used to position the modified ISG15 near the Arp3 lysine residues, K18, K75, and K191. This was done with two different Arp2/3 complexes: the inactive complex (PDB ID: 6YW7) and the active complex (PDB ID: 7AQK).

### AI analysis of *Listeria* actin comet tails

For the quantitative analysis of the actin comet tail images, we developed an image annotation tool and a custom analysis workflow to extract morphological information from the images. The image annotation tool is built on top of ImJoy (https://imjoy.io/#/, a web-based computational platform for developing interactive data analysis tools) and Kaibu (https://kaibu.org, a browser-based image annotation plugin for ImJoy). The multi-channel actin comet tail images are preprocessed to render as color images in PNG format and hosted on a server, then a sharable link was generated for remote annotation of the comet tails. During the annotation, the annotators were instructed to draw polygons around the tail for both the actin channel and the Arp3 channel. The annotation is stored in GeoJSON format on the server and processed after the annotation.

A python script was created for performing the image quantification in order to extract the mean curvature, length and area of each annotated actin comet tail. It is done by generating a label image of all the comet tail in the microscopy image. Each comet tail in the image were assigned with a unique ID (from 1 to the total number of comet tails in the image) and the resulting labeled image was generated by filling each comet tail polygon with a pixel value equals to its ID. The label image with the tails are then processed with mainly two python modules: scikit-image (https://scikit-image.org/, a Python image analysis library) and skan(https://skeleton-analysis.org/, a Python library for analyzing skeleton images). More specifically, we used skimage.measure.regionprops to compute the pixel area of each act comet tail. For computing the length and curvature of the actin comet tail, the labeled images were skeletonized with the skimage.morphology.skeletonize function. For each skeletonized tail, a line path was extracted with the member functions under each skan.Skeleton object. The path was then used to compute the mean curvature and length for each tail.

### Quantification and statistical analysis

To determine statistical significance in animal experiments, two-tailed Kaplan-Meier survival analysis was used to compare the survival percentages at each time point. For quantification of bacterial populations, measurements of actin comet tails, and in vitro cell-to-cell spread assays, we first determined whether datasets presented with normal distribution and equal standard deviations (SD) across samples. We then used one-way ANOVA followed by Tukey’s test post hoc for those with a Gaussian distribution and had equal SDs. Kruskal-Wallis test with Dunn’s multiple comparison post hoc was used when the null hypothesis was rejected. We used ANOVA with Brown-Forsythe and Welch’s correction with Dunnett’s T3 multiple comparisons post hoc when we analyzed selected pairs of comparisons with significantly different variances. We used unpaired Kolmogorov-Smirnov t-test for two-sample comparisons, when the null hypothesis was rejected. The statistical methods for the proteomics analysis are discussed in the proteomics methods of our previously published paper. Exact p-values are displayed for all statistical analyses.

## Code availability

The actin comet tail images along with the annotations will be publicly accessible via Zenodo.

The source code for performing the actin comet tail quantification is freely accessible via Github: https://github.dev/radoshevichlab/YZhang

## References

1 Perng, Y.-C. & Lenschow, D. J. in Nature Reviews Microbiology Vol. 16 423–439 (Nature Publishing Group, 2018).

2 Bogunovic, D., Byun, M., Durfee, L. A., Abhyankar, A., Sanal, O., Mansouri, D., Salem, S., Radovanovic, I., Grant, A. V., Adimi, P., Mansouri, N., Okada, S., Bryant, V. L., Kong, X.-F., Kreins, A., Velez, M. M., Boisson, B., Khalilzadeh, S., Ozcelik, U., Darazam, I. A., Schoggins, J. W., Rice, C. M., Al-Muhsen, S., Behr, M., Vogt, G., Puel, A., Bustamante, J., Gros, P., Huibregtse, J. M., Abel, L., Boisson-Dupuis, S. & Casanova, J.-L. in Science (New York, N.Y.) Vol. 337 1684–1688 (American Association for the Advancement of Science, 2012).

3 Werneke, S. W., Schilte, C., Rohatgi, A., Monte, K. J., Michault, A., Arenzana-Seisdedos, F., Vanlandingham, D. L., Higgs, S., Fontanet, A., Albert, M. L. & Lenschow, D. J. ISG15 is critical in the control of Chikungunya virus infection independent of UbE1L mediated conjugation. PLoS pathogens 7, e1002322, doi:10.1371/journal.ppat.1002322 (2011).

4 Krug, R. M., Zhao, C. & Beaudenon, S. Properties of the ISG15 E1 enzyme UbE1L. Methods Enzymol 398, 32–40, doi:10.1016/S0076-6879(05)98004-X (2005).

5 Kim, K. I., Giannakopoulos, N. V., Virgin, H. W. & Zhang, D. E. Interferon-inducible ubiquitin E2, Ubc8, is a conjugating enzyme for protein ISGylation. Molecular and cellular biology 24, 9592–9600, doi:10.1128/MCB.24.21.9592-9600.2004 (2004).

6 Durfee, L. A., Kelley, M. L. & Huibregtse, J. M. The basis for selective E1-E2 interactions in the ISG15 conjugation system. The Journal of biological chemistry 283, 23895–23902, doi:10.1074/jbc.M804069200 (2008).

7 Wong, J. J., Pung, Y. F., Sze, N. S. & Chin, K. C. HERC5 is an IFN-induced HECT-type E3 protein ligase that mediates type I IFN-induced ISGylation of protein targets. Proceedings of the National Academy of Sciences of the United States of America 103, 10735–10740, doi:10.1073/pnas.0600397103 (2006).

8 Malakhov, M. P., Malakhova, O. A., Kim, K. I., Ritchie, K. J. & Zhang, D. E. UBP43 (USP18) specifically removes ISG15 from conjugated proteins. The Journal of biological chemistry 277, 9976–9981, doi:10.1074/jbc.M109078200 (2002).

9 Zhang, X., Bogunovic, D., Payelle-Brogard, B., Francois-Newton, V., Speer, S. D., Yuan, C., Volpi, S., Li, Z., Sanal, O., Mansouri, D., Tezcan, I., Rice, G. I., Chen, C., Mansouri, N., Mahdaviani, S. A., Itan, Y., Boisson, B., Okada, S., Zeng, L., Wang, X., Jiang, H., Liu, W., Han, T., Liu, D., Ma, T., Wang, B., Liu, M., Liu, J. Y., Wang, Q. K., Yalnizoglu, D., Radoshevich, L., Uze, G., Gros, P., Rozenberg, F., Zhang, S. Y., Jouanguy, E., Bustamante, J., Garcia-Sastre, A., Abel, L., Lebon, P., Notarangelo, L. D., Crow, Y. J., Boisson-Dupuis, S., Casanova, J. L. & Pellegrini, S. Human intracellular ISG15 prevents interferon-alpha/beta over-amplification and auto-inflammation. Nature 517, 89–93, doi:10.1038/nature13801 (2015).

10 Martin-Fernandez, M., Bravo Garcia-Morato, M., Gruber, C., Murias Loza, S., Malik, M. N. H., Alsohime, F., Alakeel, A., Valdez, R., Buta, S., Buda, G., Marti, M. A., Larralde, M., Boisson, B., Feito Rodriguez, M., Qiu, X., Chrabieh, M., Al Ayed, M., Al Muhsen, S., Desai, J. V., Ferre, E. M. N., Rosenzweig, S. D., Amador-Borrero, B., Bravo-Gallego, L. Y., Olmer, R., Merkert, S., Bret, M., Sood, A. K., Al-Rabiaah, A., Temsah, M. H., Halwani, R., Hernandez, M., Pessler, F., Casanova, J. L., Bustamante, J., Lionakis, M. S. & Bogunovic, D. Systemic Type I IFN Inflammation in Human ISG15 Deficiency Leads to Necrotizing Skin Lesions. Cell Rep 31, 107633, doi:10.1016/j.celrep.2020.107633 (2020).

11 Malik, M. N. H., Waqas, S. F., Zeitvogel, J., Cheng, J., Geffers, R., Gouda, Z. A., Elsaman, A. M., Radwan, A. R., Schefzyk, M., Braubach, P., Auber, B., Olmer, R., Musken, M., Roesner, L. M., Gerold, G., Schuchardt, S., Merkert, S., Martin, U., Meissner, F., Werfel, T. & Pessler, F. Congenital deficiency reveals critical role of ISG15 in skin homeostasis. J Clin Invest 132, doi:10.1172/JCI141573 (2022).

12 Bogunovic, D., Boisson-Dupuis, S. & Casanova, J. L. ISG15: leading a double life as a secreted molecule. Exp Mol Med 45, e18, doi:10.1038/emm.2013.36 (2013).

13 Lenschow, D. J., Lai, C., Frias-Staheli, N., Giannakopoulos, N. V., Lutz, A., Wolff, T., Osiak, A., Levine, B., Schmidt, R. E., Garcia-Sastre, A., Leib, D. A., Pekosz, A., Knobeloch, K. P., Horak, I. & Virgin, H. W. t. IFN-stimulated gene 15 functions as a critical antiviral molecule against influenza, herpes, and Sindbis viruses. Proceedings of the National Academy of Sciences of the United States of America 104, 1371–1376, doi:10.1073/pnas.0607038104 (2007).

14 Lebreton, A., Lakisic, G., Job, V., Fritsch, L., Tham, T. N., Camejo, A., Mattei, P. J., Regnault, B., Nahori, M. A., Cabanes, D., Gautreau, A., Ait-Si-Ali, S., Dessen, A., Cossart, P. & Bierne, H. A bacterial protein targets the BAHD1 chromatin complex to stimulate type III interferon response. Science 331, 1319–1321, doi:10.1126/science.1200120 (2011).

15 Woodward, J. J., Iavarone, A. T. & Portnoy, D. A. c-di-AMP secreted by intracellular Listeria monocytogenes activates a host type I interferon response. Science 328, 1703–1705, doi:10.1126/science.1189801 (2010).

16 Sauer, J. D., Sotelo-Troha, K., von Moltke, J., Monroe, K. M., Rae, C. S., Brubaker, S. W., Hyodo, M., Hayakawa, Y., Woodward, J. J., Portnoy, D. A. & Vance, R. E. The N-ethyl-N-nitrosourea-induced Goldenticket mouse mutant reveals an essential function of Sting in the in vivo interferon response to Listeria monocytogenes and cyclic dinucleotides. Infect Immun 79, 688–694, doi:10.1128/IAI.00999-10 (2011).

17 Herskovits, A. A., Auerbuch, V. & Portnoy, D. A. Bacterial ligands generated in a phagosome are targets of the cytosolic innate immune system. PLoS Pathog 3, e51, doi:10.1371/journal.ppat.0030051 (2007).

18 Archer, K. A., Durack, J. & Portnoy, D. A. STING-dependent type I IFN production inhibits cell-mediated immunity to Listeria monocytogenes. PLoS Pathog 10, e1003861, doi:10.1371/journal.ppat.1003861 (2014).

19 Radoshevich, L. & Cossart, P. Listeria monocytogenes: towards a complete picture of its physiology and pathogenesis. Nat Rev Microbiol 16, 32–46, doi:10.1038/nrmicro.2017.126 (2018).

20 Auerbuch, V., Brockstedt, D. G., Meyer-Morse, N., O’Riordan, M. & Portnoy, D. A. Mice lacking the type I interferon receptor are resistant to Listeria monocytogenes. J Exp Med 200, 527–533, doi:10.1084/jem.20040976 (2004).

21 Stockinger, S., Reutterer, B., Schaljo, B., Schellack, C., Brunner, S., Materna, T., Yamamoto, M., Akira, S., Taniguchi, T., Murray, P. J., Muller, M. & Decker, T. IFN regulatory factor 3-dependent induction of type I IFNs by intracellular bacteria is mediated by a TLR- and Nod2-independent mechanism. Journal of immunology 173, 7416–7425, doi:10.4049/jimmunol.173.12.7416 (2004).

22 Carrero, J. A., Calderon, B. & Unanue, E. R. Type I interferon sensitizes lymphocytes to apoptosis and reduces resistance to Listeria infection. J Exp Med 200, 535–540, doi:10.1084/jem.20040769 (2004).

23 Radoshevich, L., Impens, F., Ribet, D., Quereda, J. J., Tham, T. N., Nahori, M. A., Bierne, H., Dussurget, O., Pizarro-Cerdá, J., Knobeloch, K. P. & Cossart, P. in eLife Vol. 4 1–23 (2015).

24 Kimmey, J. M., Campbell, J. A., Weiss, L. A., Monte, K. J., Lenschow, D. J. & Stallings, C. L. in Microbes and infection Vol. 19 249–258 (NIH Public Access, 2017).

25 Ketscher, L., Hannß, R., Morales, D. J., Basters, A., Guerra, S., Goldmann, T., Hausmann, A., Prinz, M., Naumann, R., Pekosz, A., Utermöhlen, O., Lenschow, D. J. & Knobeloch, K.-P. in Proceedings of the National Academy of Sciences of the United States of America Vol. 112 1577–1582 (National Academy of Sciences, 2015).

26 Zhang, Y., Thery, F., Wu, N. C., Luhmann, E. K., Dussurget, O., Foecke, M., Bredow, C., Jimenez-Fernandez, D., Leandro, K., Beling, A., Knobeloch, K. P., Impens, F., Cossart, P. & Radoshevich, L. The in vivo ISGylome links ISG15 to metabolic pathways and autophagy upon Listeria monocytogenes infection. Nat Commun 10, 5383, doi:10.1038/s41467-019-13393-x (2019).

27 Van Troys, M., Lambrechts, A., David, V., Demol, H., Puype, M., Pizarro-Cerda, J., Gevaert, K., Cossart, P. & Vandekerckhove, J. The actin propulsive machinery: the proteome of Listeria monocytogenes tails. Biochemical and biophysical research communications 375, 194–199, doi:10.1016/j.bbrc.2008.07.152 (2008).

28 Giannakopoulos, N. V., Luo, J. K., Papov, V., Zou, W., Lenschow, D. J., Jacobs, B. S., Borden, E. C., Li, J., Virgin, H. W. & Zhang, D. E. Proteomic identification of proteins conjugated to ISG15 in mouse and human cells. Biochemical and biophysical research communications 336, 496–506, doi:10.1016/j.bbrc.2005.08.132 (2005).

29 Becavin, C., Bouchier, C., Lechat, P., Archambaud, C., Creno, S., Gouin, E., Wu, Z., Kuhbacher, A., Brisse, S., Pucciarelli, M. G., Garcia-del Portillo, F., Hain, T., Portnoy, D. A., Chakraborty, T., Lecuit, M., Pizarro-Cerda, J., Moszer, I., Bierne, H. & Cossart, P. Comparison of widely used Listeria monocytogenes strains EGD, 10403S, and EGD-e highlights genomic variations underlying differences in pathogenicity. mBio 5, e00969–00914, doi:10.1128/mBio.00969-14 (2014).

30 Welch, M. D. & Way, M. Arp2/3-mediated actin-based motility: a tail of pathogen abuse. Cell host & microbe 14, 242–255, doi:10.1016/j.chom.2013.08.011 (2013).

31 Cudmore, S., Reckmann, I., Griffiths, G. & Way, M. Vaccinia virus: a model system for actin-membrane interactions. J Cell Sci 109 (Pt 7), 1739–1747, doi:10.1242/jcs.109.7.1739 (1996).

32 Dodding, M. P., Mitter, R., Humphries, A. C. & Way, M. A kinesin-1 binding motif in vaccinia virus that is widespread throughout the human genome. The EMBO journal 30, 4523–4538, doi:10.1038/emboj.2011.326 (2011).

33 Theriot, J. A., Mitchison, T. J., Tilney, L. G. & Portnoy, D. A. The rate of actin-based motility of intracellular Listeria monocytogenes equals the rate of actin polymerization. Nature 357, 257–260, doi:10.1038/357257a0 (1992).

34 Alberts, J. B. & Odell, G. M. In silico reconstitution of Listeria propulsion exhibits nano-saltation. PLoS Biol 2, e412, doi:10.1371/journal.pbio.0020412 (2004).

35 von Loeffelholz, O., Purkiss, A., Cao, L., Kjaer, S., Kogata, N., Romet-Lemonne, G., Way, M. & Moores, C. A. Cryo-EM of human Arp2/3 complexes provides structural insights into actin nucleation modulation by ARPC5 isoforms. Biol Open 9, doi:10.1242/bio.054304 (2020).

36 Kocks, C., Gouin, E., Tabouret, M., Berche, P., Ohayon, H. & Cossart, P. L. monocytogenes-induced actin assembly requires the actA gene product, a surface protein. Cell 68, 521–531, doi:10.1016/0092-8674(92)90188-i (1992).

37 Yoshikawa, Y., Ogawa, M., Hain, T., Chakraborty, T. & Sasakawa, C. Listeria monocytogenes ActA is a key player in evading autophagic recognition. Autophagy 5, 1220–1221 (2009).

38 Pistor, S., Grobe, L., Sechi, A. S., Domann, E., Gerstel, B., Machesky, L. M., Chakraborty, T. & Wehland, J. in Journal of cell science Vol. 113 (Pt 1 3277–3287 (J Cell Sci, 2000).

39 Weeks, B. S. & Friedman, H. M. Laminin reduces HSV-1 spread from cell to cell in human keratinocyte cultures. Biochemical and biophysical research communications 230, 466–469, doi:10.1006/bbrc.1996.5925 (1997).

40 Engelbrecht, F., Dickneite, C., Lampidis, R., Gotz, M., DasGupta, U. & Goebel, W. Sequence comparison of the chromosomal regions encompassing the internalin C genes (inlC) of Listeria monocytogenes and L. ivanovii. Mol Gen Genet 257, 186–197, doi:10.1007/s004380050638 (1998).

41 Kuhbacher, A., Emmenlauer, M., Ramo, P., Kafai, N., Dehio, C., Cossart, P. & Pizarro-Cerda, J. Genome-Wide siRNA Screen Identifies Complementary Signaling Pathways Involved in Listeria Infection and Reveals Different Actin Nucleation Mechanisms during Listeria Cell Invasion and Actin Comet Tail Formation. mBio 6, e00598–00515, doi:10.1128/mBio.00598-15 (2015).

42 Ortega, F. E., Koslover, E. F. & Theriot, J. A. Listeria monocytogenes cell-to-cell spread in epithelia is heterogeneous and dominated by rare pioneer bacteria. eLife 8, doi:10.7554/eLife.40032 (2019).

43 Siegrist, M. S., Aditham, A. K., Espaillat, A., Cameron, T. A., Whiteside, S. A., Cava, F., Portnoy, D. A. & Bertozzi, C. R. Host actin polymerization tunes the cell division cycle of an intracellular pathogen. Cell Rep 11, 499–507, doi:10.1016/j.celrep.2015.03.046 (2015).

44 Pizarro-Cerdá, J., Chorev, D. S., Geiger, B. & Cossart, P. in Trends in cell biology Vol. 27 93–100 (Elsevier Current Trends, 2017).

45 Swaim, C. D., Scott, A. F., Canadeo, L. A. & Huibregtse, J. M. Extracellular ISG15 Signals Cytokine Secretion through the LFA-1 Integrin Receptor. Molecular cell 68, 581–590 e585, doi:10.1016/j.molcel.2017.10.003 (2017).

46 Sridharan, H., Zhao, C. & Krug, R. M. Species specificity of the NS1 protein of influenza B virus: NS1 binds only human and non-human primate ubiquitin-like ISG15 proteins. The Journal of biological chemistry 285, 7852–7856, doi:10.1074/jbc.C109.095703 (2010).

47 Durfee, L. A., Lyon, N., Seo, K. & Huibregtse, J. M. The ISG15 conjugation system broadly targets newly synthesized proteins: implications for the antiviral function of ISG15. Molecular cell 38, 722–732, doi:10.1016/j.molcel.2010.05.002 (2010).

48 Swaim, C. D., Canadeo, L. A., Monte, K. J., Khanna, S., Lenschow, D. J. & Huibregtse, J. M. Modulation of Extracellular ISG15 Signaling by Pathogens and Viral Effector Proteins. Cell Rep 31, 107772, doi:10.1016/j.celrep.2020.107772 (2020).

49 Munnur, D., Teo, Q., Eggermont, D., Lee, H. H. Y., Thery, F., Ho, J., van Leur, S. W., Ng, W. W. S., Siu, L. Y. L., Beling, A., Ploegh, H., Pinto-Fernandez, A., Damianou, A., Kessler, B., Impens, F., Mok, C. K. P. & Sanyal, S. Altered ISGylation drives aberrant macrophage-dependent immune responses during SARS-CoV-2 infection. Nat Immunol 22, 1416–1427, doi:10.1038/s41590-021-01035-8 (2021).

50 Speer, S. D., Li, Z., Buta, S., Payelle-Brogard, B., Qian, L., Vigant, F., Rubino, E., Gardner, T. J., Wedeking, T., Hermann, M., Duehr, J., Sanal, O., Tezcan, I., Mansouri, N., Tabarsi, P., Mansouri, D., Francois-Newton, V., Daussy, C. F., Rodriguez, M. R., Lenschow, D. J., Freiberg, A. N., Tortorella, D., Piehler, J., Lee, B., Garcia-Sastre, A., Pellegrini, S. & Bogunovic, D. ISG15 deficiency and increased viral resistance in humans but not mice. Nat Commun 7, 11496, doi:10.1038/ncomms11496 (2016).

51 Waqas, S. F., Sohail, A., Nguyen, A. H. H., Usman, A., Ludwig, T., Wegner, A., Malik, M. N. H., Schuchardt, S., Geffers, R., Winterhoff, M., Merkert, S., Martin, U., Olmer, R., Lachmann, N. & Pessler, F. ISG15 deficiency features a complex cellular phenotype that responds to treatment with itaconate and derivatives. Clin Transl Med 12, e931, doi:10.1002/ctm2.931 (2022).

52 Stockinger, S., Materna, T., Stoiber, D., Bayr, L., Steinborn, R., Kolbe, T., Unger, H., Chakraborty, T., Levy, D. E., Muller, M. & Decker, T. Production of type I IFN sensitizes macrophages to cell death induced by Listeria monocytogenes. Journal of immunology 169, 6522–6529, doi:10.4049/jimmunol.169.11.6522 (2002).

53 Rayamajhi, M., Humann, J., Penheiter, K., Andreasen, K. & Lenz, L. L. Induction of IFN-alphabeta enables Listeria monocytogenes to suppress macrophage activation by IFN-gamma. J Exp Med 207, 327–337, doi:10.1084/jem.20091746 (2010).

54 Osborne, S. E., Sit, B., Shaker, A., Currie, E., Tan, J. M. J., van Rijn, J., Higgins, D. E. & Brumell, J. H. in Cellular microbiology Vol. 19 e12660 (John Wiley & Sons, Ltd, 2017).

55 Tan, J. M. J., Garner, M. E., Regeimbal, J. M., Greene, C. J., Marquez, J. D. R., Ammendolia, D. A., McCluggage, A. R. R., Li, T., Wu, K. J., Cemma, M., Ostrowski, P. P., Raught, B., Diamond, M. S., Grinstein, S., Yates, R. M., Higgins, D. E. & Brumell, J. H. Listeria exploits IFITM3 to suppress antibacterial activity in phagocytes. Nat Commun 12, 4999, doi:10.1038/s41467-021-24982-0 (2021).

56 Phuyal, S., Suarez, S. S. & Tung, C. K. Biological benefits of collective swimming of sperm in a viscoelastic fluid. Front Cell Dev Biol 10, 961623, doi:10.3389/fcell.2022.961623 (2022).

57 Mostowy, S. & Shenoy, A. R. in Nature reviews. Immunology Vol. 15 559–573 (Europe PMC Funders, 2015).

58 Wells, A. I. & Coyne, C. B. Type III Interferons in Antiviral Defenses at Barrier Surfaces. Trends Immunol 39, 848–858, doi:10.1016/j.it.2018.08.008 (2018).

59 Rusinova, I., Forster, S., Yu, S., Kannan, A., Masse, M., Cumming, H., Chapman, R. & Hertzog, P. J. Interferome v2.0: an updated database of annotated interferon-regulated genes. Nucleic Acids Res 41, D1040–1046, doi:10.1093/nar/gks1215 (2013).

60 Zuo, C., Sheng, X., Ma, M., Xia, M. & Ouyang, L. ISG15 in the tumorigenesis and treatment of cancer: An emerging role in malignancies of the digestive system. Oncotarget 7, 74393–74409, doi:10.18632/oncotarget.11911 (2016).

61 Timmer, T., Terpstra, P., van den Berg, A., Veldhuis, P. M., Ter Elst, A., van der Veen, A. Y., Kok, K., Naylor, S. L. & Buys, C. H. An evolutionary rearrangement of the Xp11.3-11.23 region in 3p21.3, a region frequently deleted in a variety of cancers. Genomics 60, 238–240, doi:10.1006/geno.1999.5878 (1999).

62 Desai, S. D., Reed, R. E., Burks, J., Wood, L. M., Pullikuth, A. K., Haas, A. L., Liu, L. F., Breslin, J. W., Meiners, S. & Sankar, S. ISG15 disrupts cytoskeletal architecture and promotes motility in human breast cancer cells. Experimental biology and medicine 237, 38–49, doi:10.1258/ebm.2011.011236 (2012).

63 Song, F., Zhang, Y., Pan, Z., Hu, X., Yi, Y., Zheng, X., Wei, H. & Huang, P. in Cancer Cell International Vol. 21 1–16 (BioMed Central Ltd, 2021).

## References for Methods

1 Li, F., Adase, C. A. & Zhang, L. J. Isolation and Culture of Primary Mouse Keratinocytes from Neonatal and Adult Mouse Skin. J Vis Exp, doi:10.3791/56027 (2017).

2 Lauer, P., Chow, M. Y., Loessner, M. J., Portnoy, D. A. & Calendar, R. Construction, characterization, and use of two Listeria monocytogenes site-specific phage integration vectors. J Bacteriol 184, 4177–4186, doi:10.1128/JB.184.15.4177-4186.2002 (2002).

3 Balestrino, D., Anne Hamon, M., Dortet, L., Nahori, M. A., Pizarro-Cerda, J., Alignani, D., Dussurget, O., Cossart, P. & Toledo-Arana, A. in Applied and environmental microbiology Vol. 76 3625 (American Society for Microbiology (ASM), 2010).

4 Morgenstern, J. P. & Land, H. Advanced mammalian gene transfer: high titre retroviral vectors with multiple drug selection markers and a complementary helper-free packaging cell line. Nucleic Acids Res 18, 3587–3596, doi:10.1093/nar/18.12.3587 (1990).

5 Cong, L., Ran, F. A., Cox, D., Lin, S., Barretto, R., Habib, N., Hsu, P. D., Wu, X., Jiang, W., Marraffini, L. A. & Zhang, F. Multiplex genome engineering using CRISPR/Cas systems. Science 339, 819–823, doi:10.1126/science.1231143 (2013).

6 Ran, F. A., Hsu, P. D., Wright, J., Agarwala, V., Scott, D. A. & Zhang, F. Genome engineering using the CRISPR-Cas9 system. Nat Protoc 8, 2281–2308, doi:10.1038/nprot.2013.143 (2013).

7 Cacciabue, M., Currá, A. & Gismondi, M. I. ViralPlaque: a Fiji macro for automated assessment of viral plaque statistics. PeerJ 7, e7729, doi:10.7717/peerj.7729 (2019).

8 Kurup, S. P., Obeng-Adjei, N., Anthony, S. M., Traore, B., Doumbo, O. K., Butler, N. S., Crompton, P. D. & Harty, J. T. Regulatory T cells impede acute and long-term immunity to blood-stage malaria through CTLA-4. Nat Med 23, 1220–1225, doi:10.1038/nm.4395 (2017).

9 Meyerholz, D. K. & Beck, A. P. Principles and approaches for reproducible scoring of tissue stains in research. Lab Invest 98, 844–855, doi:10.1038/s41374-018-0057-0 (2018).

10 Meijering, E., Dzyubachyk, O. & Smal, I. Methods for cell and particle tracking. Methods Enzymol 504, 183–200, doi:10.1016/b978-0-12-391857-4.00009-4 (2012).

11 Stirling, D. R., Swain-Bowden, M. J., Lucas, A. M., Carpenter, A. E., Cimini, B. A. & Goodman, A. CellProfiler 4: improvements in speed, utility and usability. BMC Bioinformatics 22, 433, doi:10.1186/s12859-021-04344-9 (2021).

12 Logan, D. J., Shan, J., Bhatia, S. N. & Carpenter, A. E. Quantifying co-cultured cell phenotypes in high-throughput using pixel-based classification. Methods 96, 6–11, doi:10.1016/j.ymeth.2015.12.002 (2016).

13 Zhang, Y., Thery, F., Wu, N. C., Luhmann, E. K., Dussurget, O., Foecke, M., Bredow, C., Jimenez-Fernandez, D., Leandro, K., Beling, A., Knobeloch, K. P., Impens, F., Cossart, P. & Radoshevich, L. The in vivo ISGylome links ISG15 to metabolic pathways and autophagy upon Listeria monocytogenes infection. Nat Commun 10, 5383, doi:10.1038/s41467-019-13393-x (2019).

14 Van Troys, M., Lambrechts, A., David, V., Demol, H., Puype, M., Pizarro-Cerda, J., Gevaert, K., Cossart, P. & Vandekerckhove, J. The actin propulsive machinery: the proteome of Listeria monocytogenes tails. Biochemical and biophysical research communications 375, 194–199, doi:10.1016/j.bbrc.2008.07.152 (2008).

